# Interplay between the cyclophilin homology domain of RANBP2 and MX2 regulates HIV-1 capsid dependencies on nucleoporins

**DOI:** 10.1101/2024.08.12.607676

**Authors:** Haley Flick, Ananya Venbakkam, Parmit K. Singh, Bailey Layish, Szu-Wei Huang, Rajalingam Radhakrishnan, Mamuka Kvaratskhelia, Alan N. Engelman, Melissa Kane

## Abstract

Interlinked interactions between the viral capsid (CA), nucleoporins (Nups), the antiviral protein myxovirus resistance 2 (MX2/MXB) influence HIV-1 nuclear entry and the outcome of infection. Although RANBP2/NUP358 has been repeatedly identified as a critical player in HIV-1 nuclear import and MX2 activity, the mechanism by which RANBP2 facilitates HIV-1 infection is not well understood. To explore the interactions between MX2, the viral CA, and RANBP2, we utilized CRISPR-Cas9 to generate cell lines expressing RANBP2 from its endogenous locus but lacking the C-terminal cyclophilin (Cyp) homology domain, and found that both HIV-1 and HIV-2 infection were reduced significantly in RANBP2_ΔCyp_ cells. Importantly, although MX2 still localized to the nuclear pore complex in RANBP2_ΔCyp_ cells, antiviral activity against HIV-1 was decreased. By generating cells expressing specific point mutations in the RANBP2-Cyp domain we determined that the effect of the RANBP2-Cyp domain on MX2 anti-HIV-1 activity is due to direct interactions between RANBP2 and CA. We further determined that CypA and RANBP2-Cyp have similar effects on HIV-1 integration targeting. Finally, we found that the Nup requirements for HIV infection and MX2 activity were altered in cells lacking the RANBP2-Cyp domain. These findings demonstrate that the RANBP2-Cyp domain affects viral infection and MX2 sensitivity by altering CA-specific interactions with cellular factors that affect nuclear import and integration targeting.

**Significance Statement:** HIV-1 entry into the nucleus is an essential step in viral replication that involves complex interactions between the viral capsid and multiple cellular proteins, including nucleoporins such as RANBP2. Nups also mediate the function of the antiviral protein MX2, however determining the precise role of Nups in HIV infection has proved challenging due to the complex nature of the nuclear pore and significant pleiotropic effects elicited by Nup depletion. We have used precise gene editing to assess the role of the Cyp domain of RANBP2 in HIV-1 infection and MX2 activity. We find that this domain affects viral infection, nucleoporin requirements, MX2 sensitivity, and integration targeting in a CA-specific manner, providing detailed insights into how RANBP2 contributes to HIV-1 infection.

## Introduction

Access to the chromosomal DNA contained within the nucleus of target cells is critical for retroviral integration and replication. Among retroviruses, the lentiviruses are uniquely efficient in their ability to enter the nucleus of interphase cells, in which the nuclear membrane is intact. The viral capsid (CA) is the key viral determinant of the ability of HIV-1 to infect non-dividing cells (1) and recent research has indicated that the mature capsid lattice mimics cellular nuclear transport receptors (NTRs) to mediate high valency interactions with FG-nucleoporins (Nups) that compose the inner channel of the nuclear pore complex (NPC) (2, 3). Other CA-interacting host proteins, including the peptidylprolyl isomerase cyclophilin A (CypA) (4, 5) and the mRNA processing protein cleavage and polyadenylation specificity factor 6 (CPSF6) (6), can also influence HIV-1 nuclear transport, perhaps via regulating CA-Nup interactions (7, 8) CA-binding host factors implicated in viral nuclear import can furthermore influence sites of HIV-1 integration in the human genome perhaps by directing preintegration complexes to specific nuclear import pathways, by affecting downstream interactions with CPSF6, which is a key regulator of viral nuclear incursion (9), or through effects on chromatin architecture (10).

RANBP2(NUP358) is a metazoan-specific component of the cytoplasmic face component of the NPC. The 3224 residue human protein is the largest component of the NPC and is anchored to the cytoplasmic outer ring via its N-terminal domain (NTD) in pentameric bundles connected by interactions between oligomerization elements (OE) [Figure 1A and (11, 12)]. The remaining domains, including several FG repeats, four Ran-binding domains, and a Cyp homology domain, extend into the cytoplasm and are connected by unstructured linker sequences (11, 13). Numerous reports using RNAi-mediated depletion have indicated that RANBP2 plays a major role in nuclear import of HIV-1 and HIV-2 (5, 14–24); however, due to the importance of RANBP2 in multiple cellular processes (25–27), and the numerous pleiotropic effects that can occur upon Nup depletion (18), these results must be interpreted with caution. The C-terminal Cyp domain of RANBP2 contains peptidyl-prolyl isomerase activity, and both genetic and biochemical studies have demonstrated that it directly interacts with HIV-1 CA (5, 15, 21, 22, 28, 29). Furthermore, HIV-1 CA mutant viruses G89V and P90A, which are impaired for CypA binding, exhibit reduced dependence on RANBP2 for infection (5, 18). However, the precise role of the RANBP2 Cyp domain in HIV-1 infection remains controversial. On the one hand, some reports have proposed that CA-RANBP2 interactions are crucial for infection and are a key driver of species-specific primate lentiviral adaptation (21, 29). On the other hand, experiments utilizing ectopic expression of truncated human RANBP2 in mouse cells suggested that direct interactions with the RANBP2-Cyp domain are not required for HIV-1 nuclear import (22).

**Fig. 1.**
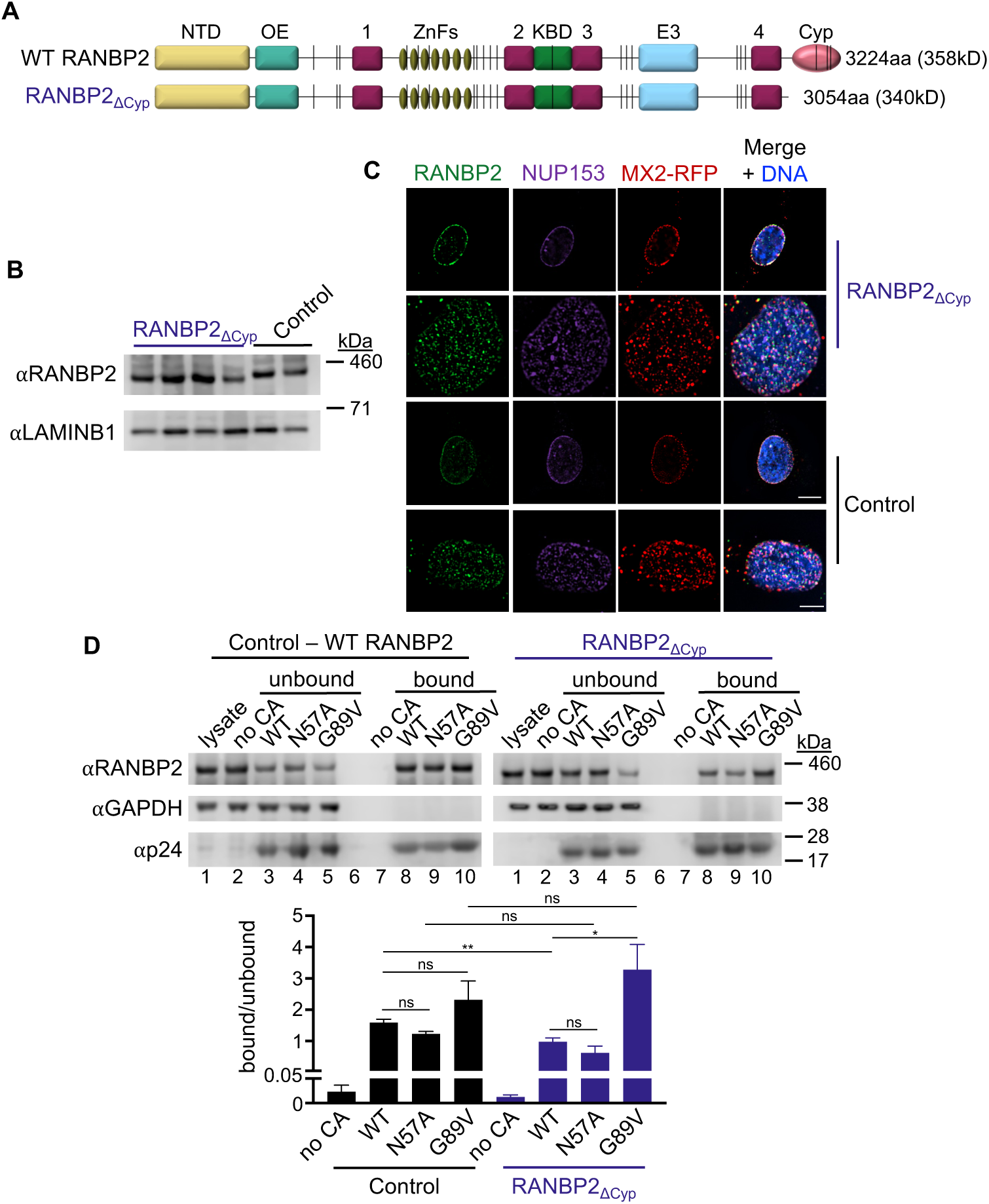
The RANBP2-Cyp domain is not required for MX2 localization to the nuclear envelope or for HIV-1 CA binding. (*A*) Domain architecture of WT RANBP2 and RANBP2_ΔCyp_. NTD (yellow), N-terminal domain; OE (cyan) oligomerization element; 1-4 (magenta), Ran binding domains; ZnFs (gold), zinc finger motifs; KBD (green), kinesin-binding domain; E3 (blue), SUMO E3 ligase domain; Cyp (pink), cyclophilin homology domain; vertical lines (black) FxFG repeat. (*B*) Western blot analysis of RANBP2 expression in control and mutant cell clones and LAMINB1 loading control. (*C*) Deconvolution microscopic images (single optical sections) of control and RANBP2_ΔCyp_ cells expressing MX2-RFP (red, stably transduced with a doxycycline-inducible vector) were fixed and immunofluorescently stained for RANBP2 (green) and NUP153 (purple), and Hoescht stained for DNA. (Top) Optical sections are approximately through the center of the vertical dimension on the nucleus, scale bar = 10μm. (Bottom) – Optical sections are approximately coincident with the dorsal surface of the nucleus, scale bar = 5μm. Representative of nine RANBP2_ΔCyp_ and three control clones. (*D*) HIV-1 WT, N57A, or G89V CA tubes were assembled *in vitro* and incubated with lysates from control or RANBP2_ΔCyp_ HT1080 cells as indicated. The reaction mixtures were subjected to centrifugation to separate pulldown (bound) fractions from unbound proteins in supernatants. Western blot analysis (top) - Lane 1: cellular lysates; Lane 2: supernatant from control experiments without CA tubes; Lanes 3-5: supernatants after incubating cellular lysates with CA tubes; Lane 6: empty; Lane 7: pulled-down fraction from control experiment in the absence of CA tubes; Lanes 7-10: proteins bound to CA tubes. Blot from one RANBP2_ΔCyp_ clone of two tested is shown. Quantification (bottom): ratio of bound to unbound RANBP2 from four replicate blots of control clone and six replicate blots of two RANBP2_ΔCyp_ clones. Statistical significance was determined by one-way ANOVA (Kruskal-Wallis test); ns, not significant (*P*≥0.05); * *P* < 0.05; ** *P* < 0.01.

MX2 is a dynamin-like guanosine triphosphatase (GTPase) whose expression is strongly upregulated by type I interferons (IFNs) and localizes to the NPC via a nuclear localization-like sequence in its first 25 amino acids (30, 31). MX2 inhibits HIV-1 infection prior to chromosomal DNA integration, but after the completion of reverse transcription (30, 32, 33), and current models suggest that it acts by preventing nuclear import of the preintegration complex. HIV-1 CA is the major viral determinant of MX2 sensitivity, and several single amino-acid substitutions in CA have been identified that confer partial or complete resistance to MX2 (30, 32, 34–36). MX2 has also been found to directly bind CA, however the relevance of this binding for viral inhibition is unclear since MX2-resistant CA proteins are still efficiently bound by MX2 (37, 38). Sensitivity of HIV-1 to MX2 activity is affected by complex interactions between CA and cellular proteins involved in the early stages of HIV-1 infection [reviewed in (39)], including RANBP2. Knockdown of RANBP2 reduces the antiviral activity of MX2 against HIV-1 in multiple cell types (18, 40). MX2 is also mislocalized upon RANBP2 depletion and exhibits similar Nup requirements for recruitment to the NPC as RANBP2 (18). Additionally, localization of MX2 to the NPC following mitosis occurs subsequent to the recruitment of RANBP2 (18), which is a late event in post-mitotic NPC assembly, and follows the establishment of the structural pore and the central channel (reviewed in (41, 42), suggesting that MX2 recruitment to the NPC may be dependent on RANBP2. Finally, a yeast-two-hybrid screen indicated that RANBP2 may directly interact with the N-terminal domain of MX2, although these results were not confirmed via biochemical tests such as co-immunoprecipitation (40). The antiviral activity of MX2 is also affected by HIV-1 CA-CypA interactions, with multiple reports indicating that wild-type HIV-1 (HIV-1_WT_) is only sensitive to MX2 when it is bound to CypA (18, 35, 43, 44). CypA may affect MX2 sensitivity either by affecting viral utilization of cellular nuclear import pathways, or by producing an MX2 sensitive CA conformation or state (5, 18, 43).

In this report, we investigated the role of the Cyp domain of RANBP2 in HIV infection and MX2 activity by gene editing of the endogenous *RANBP2* locus. We find that, while abrogation of CA-Cyp domain interactions does not recapitulate the effects of RANBP2 depletion, the Cyp domain is required for efficient HIV infection and integration site selection. We also demonstrate that abrogation of RANBP2-Cyp-CA interactions reduces MX2 activity against HIV-1, but not HIV-2, and that the effects of RANBP2-Cyp on MX2 activity are mediated by direct interactions between HIV-1 CA and the Cyp domain. Finally, we show that deletion of the RANBP2-Cyp domain alters sensitivity of HIV to the depletion of other nucleoporins.

## Results

### Effects of the RANBP2-Cyp domain on MX2 localization and RANBP2-CA interactions

To generate mutant cells endogenously expressing truncated RANBP2 lacking the Cyp domain, we employed a CRISPR/Cas9-adeno associated virus (AAV)-based gene engineering approach to target the *RANBP2* locus (*SI Appendix*, Fig. S1A). We introduced Cas9 ribonucleoproteins into HT1080 cells that contained guide RNAs targeting exon 27 and exon 29 and an AAV containing donor-DNA for homology-directed repair to fuse exons 28 and 29, resulting in the deletion of the C-terminal 170 amino acids of RANBP2 (Figs. 1A and *SI Appendix*, Fig. S1A). Single cell clones with the desired biallelic mutations were confirmed by western blot (Fig. 1B) and by sequencing of both genomic DNA and mRNA. Since RANBP2 is recruited comparatively late during NPC assembly [reviewed in (41, 42)] and is attached to the cytoplasmic outer-ring spoke via its NTD (11), we would not anticipate deletion of the Cyp domain to alter nuclear envelope localization of RANBP2 or other Nups. As expected, both RANBP2 and NUP153 (which is recruited early in NPC formation [reviewed in (41, 42)]) were similarly localized in both control and RANBP2_ΔCyp_ cells. We also found that localization of an MX2-tagRFP fusion to the nuclear envelope was unaffected in RANBP2_ΔCyp_ cells (Fig. 1C) indicating that its recruitment to the NPC is independent of the RANBP2-Cyp domain.

We then tested the ability of RANBP2 in control and mutant clones to bind HIV-1 CA *in vitro* (Fig. 1D) and found that both full-length RANBP2 and RANBP2_ΔCyp_ pelleted with WT, FG-binding mutant N57A, and cyclophilin-binding mutant G89V CAs. The ratio of bound to unbound RANBP2 was similar between WT, N57A, and G89V CA, however a higher fraction of RANBP2_ΔCyp_ was bound to the G89V than WT or N57A CA. Additionally, the fraction of bound RANBP2_ΔCyp_ was slightly lower than full-length RANBP2 for the WT, but not for G89V CA. Consistent with the results of an earlier study (17), our results suggest there are both Cyp-domain-dependent and -independent interactions between RANBP2 and HIV-1 CA. Because the N57A mutant also bound RANBP2_ΔCyp_, our data indicate that Cyp-domain-independent interactions between RANBP2 and CA are unlikely to be mediated by FG motifs. Such interactions could be mediated by an unknown CA-binding motif in RANBP2 or may reflect an indirect interaction whereby RANBP2 is pelleting with CA via interactions with other host factors.

### Deletion of the RANBP2-Cyp domain affects HIV infection and MX2 sensitivity

To investigate the role of the RANBP2-Cyp domain in HIV-1 infection and MX2 sensitivity, RANBP2_ΔCyp_ cell lines were transduced with a lentiviral vector for doxycycline-inducible expression of MX2 and infected with GFP-reporter viruses in the presence or absence of the CypA inhibitor cyclosporine A (CsA) at the time of infection (Fig. 2). We observed a modest though significant reduction in HIV-1_WT_ infection (∼2.3-fold) as well as a reduction in MX2 sensitivity in RANBP2_ΔCyp_ cells (Fig. 2A), suggesting that in addition to CypA, CA-RANBP2-Cyp interactions affect MX2 activity against HIV-1. However, CsA treatment, which blocks CA-CypA, but not CA-RANBP2-Cyp interactions (5), had similar effects on sensitivity of HIV-1_WT_ to MX2 in both control and RANBP2_ΔCyp_ cells, indicating that RANBP2-Cyp may only affect MX2 activity against CAs that are already bound by CypA. Interestingly, while deletion of RANBP2-Cyp had marginal effects on infection by HIV-1_G89V CA_ as expected, the ability of MX2 to enhance infection of this mutant was reduced in RANBP2_ΔCyp_ cells (Fig. 2B), suggesting there may also be CA-independent effects of the Cyp domain on MX2 activity.

**Fig. 2.**
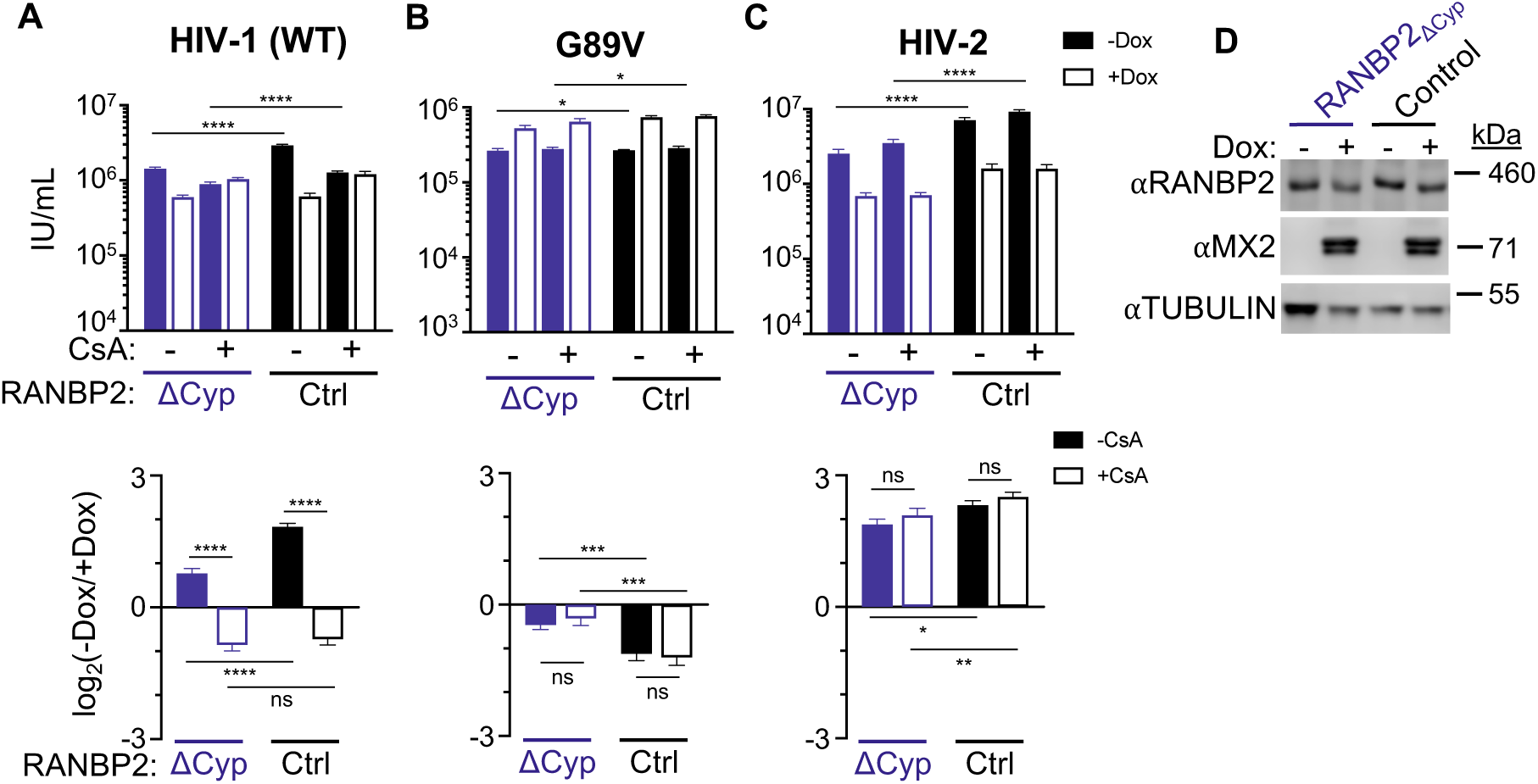
Effects of the RanBP2-Cyp domain on lentiviral infection and Mx2 sensitivity. (*A-C*) (Top) Infection of control and RANBP2_ΔCyp_ HT1080 cell clones stably transduced with doxycycline-inducible MX2 in the presence (open bars) or absence (filled bars) of doxycycline and presence or absence of CsA with the indicated GFP reporter viruses. Titers are represented as mean + sem of infectious units (IU) per mL. RANBP2_ΔCyp_: n≥12 technical replicates combined from ≥3 different clones; control: n≥8 technical replicates combined from ≥2 different clones; representative of ≥4 independent experiments. Statistical significance was determined by two-way ANOVA (Śídák’s multiple comparisons test). (Bottom) Data from (top) shown as a ratio [fold change of -Dox(-MX2)/+Dox(+MX2)] in the presence (open bars) or absence (filled bars) of CsA. Average fold change calculated from three to four technical replicates per experiment; shown is mean + sem of log_2_(fold change) from four-eight independent experiments. Statistical significance was determined by paired *t* test. D) Western blot analysis of RANBP2, doxycycline-inducible MX2, and tubulin loading control from representative clones. ns, not significant (*P*≥0.05); * *P* < 0.05; ** *P* < 0.01; *** *P* < 0.001; **** *P* < 0.0001.

In addition to HIV-1_G89V CA_, we examined the effects of RANBP2-Cyp deletion on several HIV-1 CA mutants with alterations in MX2 and CypA/CsA sensitivity, and/or impaired Nup/CPSF6 binding (*SI Appendix,* Fig. S2). Both HIV-1_N57A CA_ and the CPSF6 binding mutant N74D bind cyclophilins and are affected by CsA treatment in HT1080 cells (5, 44), but have significantly lower affinity for RANBP2-Cyp than HIV-1_WT_ (5). We found that both infectivity and MX2 sensitivity of these CA mutants was unaffected in RANBP2_ΔCyp_ cells (*SI Appendix,* Fig. S2A-B). This is also consistent with previous reports indicating that HIV-1_N57A CA_ and HIV-1_N74D CA_ are less sensitive to RANBP2 depletion (5, 20). Collectively, these findings suggest that HIV-1_N57A CA_ and HIV-1_N74D CA_ interact functionally with CypA, but not RANBP2-Cyp. The A92E and G94D mutations appear to stabilize the CypA binding loop of HIV-1 CA in a conformation resembling the CypA bound state (45), and these mutants have a complex phenotype, exhibiting CsA-resistance or -dependence and MX2-resistance that is dependent on the target cell type [reviewed in (46)]. HIV-1_A92E CA_ and HIV-1_G94D CA_ were less sensitive to RANBP2-Cyp depletion than HIV-1_WT_ (∼1.3 and ∼1.7-fold, respectively), and the sensitivity of HIV-1_G94D CA_, but not HIV-1_A92E CA_ to MX2, was modestly reduced in RANBP2_ΔCyp_ cells (*SI Appendix,* Fig. S2C-D). These findings suggest that stabilization of the CypA binding loop reduces the dependence on interactions with RANBP2-Cyp. Finally, we tested effects of RANBP2-Cyp deletion on the T210K CA mutant, which acquired MX2 resistance upon passage in MT4 cells (34), but is sensitive to MX2 in certain contexts, including upon CsA treatment (18, 44). HIV-1_T210K CA_ also has a significant infectivity defect, and has altered nucleoporin dependencies, most notably a reduced sensitivity to NUP153 depletion (18), which may be the result of both alterations in cofactor binding as well as an increase in capsid stability (47). We found that HIV-1_T210K CA_ was more sensitive than WT virus to deletion of RANBP2-Cyp (∼3.8 fold), although its sensitivity to MX2 was unaltered (*SI Appendix,* Fig. S2E). The T210K mutation alters the tri-hexamer interface of the CA lattice and affects interactions between the NTD of MX2 and the CA (47), however, our data indicate that in certain contexts, this reduced binding is sufficient to inhibit infection. Collectively, these data indicate that the RANBP2-Cyp domain plays a comparatively minor role in HIV infectivity but affects MX2 sensitivity in a viral CA sequence-dependent manner. While the full antiviral activity of MX2 against HIV-1_WT CA_ requires both CA-CypA and CA-RANBP2-Cyp interactions, this requirement is highly dependent on the CA sequence, as MX2 sensitivity of several HIV-1 CA mutants is altered by abrogation of interactions with CypA, but not RANBP2-Cyp.

### Distinct effects of RANBP2-Cyp deletion on other lentiviruses

We next explored the requirements for RANBP2-Cyp for infection of other lentiviruses with distinct MX2 sensitivities from HIV-1. While HIV-1 is of chimpanzee origin, HIV-2 (and SIVmac) arose from SIV circulating in sooty mangebeys (SIVsmm) [reviewed in (48)]. Like HIV-1, infectivity of HIV-2 and SIVmac is reduced upon RANBP2 and NUP153 depletion, however these viruses have altered interactions with a number of cellular cofactors and distinct requirements for other nucleoporins (18, 21, 49). HIV-2 also has a divergent CypA binding loop, resulting in lower affinity binding to CypA than HIV-1 (50), and inhibition of HIV-2 by MX2 is independent of CA-CypA interactions (18, 44). HIV-2 is restricted by chimeric TRIM-RANBP2-Cyp (21), indicating that it can interact with RANBP2-Cyp, however the affinity of this interaction is unknown. We found that HIV-2 was more sensitive to depletion of RANBP2-Cyp than HIV-1 (∼4.4-fold infection defect), however MX2 sensitivity of HIV-2 was only slightly reduced in RANBP2_ΔCyp_ cells (Fig. 2C). SIVmac does not bind cyclophilins (5, 51, 52), and accordingly, was not affected by deletion of RANBP2-Cyp (*SI Appendix*, Fig 3A). SIV from tantalus monkeys (SIVagmTAN) also binds cyclophilins (53–55), but it is unknown whether it requires RANBP2 for infection. We found that infectivity and MX2 sensitivity of SIVagmTAN were unaffected in RANBP2_ΔCyp_ cells (or by CsA treatment) (*SI Appendix*, Fig 3B), indicating that, at least in human cells, infectivity of HIV-2, SIVmac, and SIVagmTAN is independent of CA-Cyp interactions. We also investigated how deletion of RANBP2-Cyp affected MX2-resistant non-primate lentiviruses equine infectious anemia virus (EIAV) and feline immunodeficiency virus (FIV) (*SI Appendix*, Fig 3C-D) (30, 32). EIAV, which does not bind cyclophilins, was unaffected by RANBP2-Cyp deletion, while FIV infection was slightly elevated (∼1.3 fold) in RANBP2_ΔCyp_ cells, in agreement with previous reports indicating that RANBP2 is dispensable for EIAV and FIV infection in human cells (15, 18). These data indicate that while interactions with RANBP2-Cyp are important for HIV-2 infection, they are dispensable for infectivity of multiple other lentiviruses in human cells. However, CA-RANBP2- Cyp interactions may be involved in infection of the natural host species’ of these viruses, especially given that the RANBP2-Cyp domain exhibits evidence of evolution under positive selection (5, 56).

**Fig. 3.**
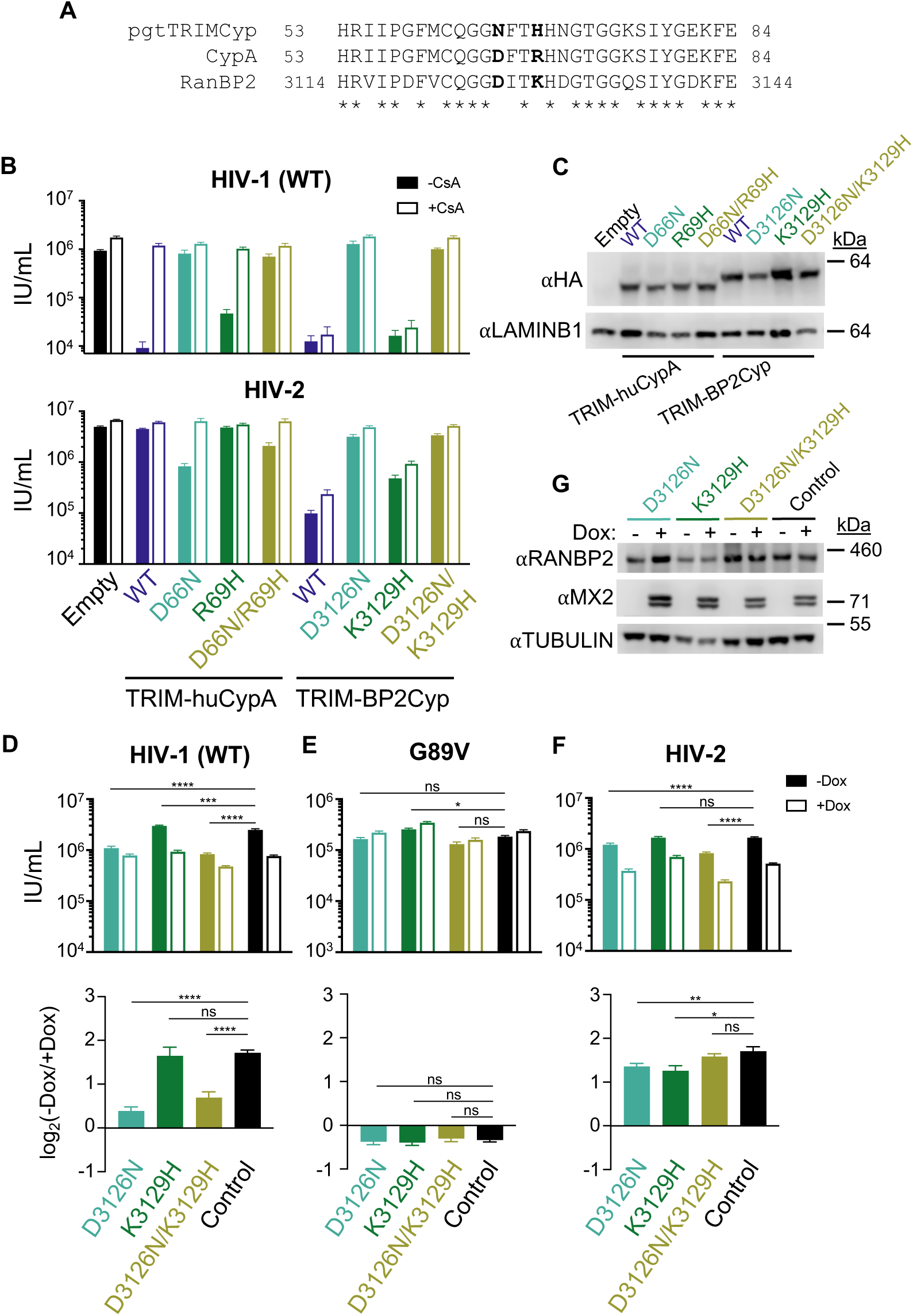
Effect of RANBP2-Cyp point mutations on lentiviral infection and MX2 sensitivity. (*A*) Amino acid alignment of portions of the Cyp domain of pig tailed/rhesus macaque (pgt) TRIMCyp, CypA, and RANBP2-Cyp with key residues involved in CA binding in bold. (*B*) Infectivity of GFP reporter viruses on HeLa cells stably expressing control empty vector, or chimeras of the TRIM5 N-terminal domain of owl monkey TRIMCyp with human CypA (huCypA), human CypA_D66N_, CypA_R69H_, CypA_D66N/R69H_ mutants, human RANBP2-Cyp (BP2-Cyp), or BP2-Cyp_D3126N_, BP2-Cyp_K3129H_, BP2-Cyp_D3126N/K3129H_ mutants. Titers are represented as mean + sem of infectious units (IU) per mL, n≥12 technical replicates combined from ≥3 independent experiments. Statistical analysis in Dataset S1. (*C*) Western blot analysis of HA-tagged TRIM fusion protein expression and LAMINB1 loading control in stable cell lines in *B*. (*D-F*) (Top) Infection of control and RANBP2 point mutant HT1080 cell clones stably transduced with doxycycline-inducible MX2 the presence (open bars) or absence (filled bars) of doxycycline with the indicated GFP reporter viruses. Titers are represented as mean + sem of infectious units (IU) per mL. n≥12 technical replicates combined from ≥4 independent experiments. Statistical significance was determined by two-way ANOVA (Śídák’s multiple comparisons test). (Bottom) Data from (top) shown as a ratio [fold change of -Dox(-MX2)/+Dox(+MX2)] in the presence (open bars) or absence (filled bars) of CsA. Average fold change calculated from three-four technical replicates per experiment; shown is mean + sem of log_2_(fold change) from four-eight independent experiments. Statistical significance was determined by paired *t* test. (*G*) Western blot analysis of RANBP2, doxycycline-inducible MX2, and tubulin loading control from clones shown in *D-F*. ns, not significant (*P*≥0.05); * *P* < 0.05; ** *P* < 0.01; *** *P* < 0.001; **** *P* < 0.0001.

### Direct interactions between RANBP2-Cyp and the HIV-1 capsid determine MX2 sensitivity

To examine the potential role of capsid-independent effects of RANBP2-Cyp on HIV-1 infection and MX2 activity, we next generated cells expressing RANBP2 from its endogenous locus but with specific point mutations introduced to the Cyp domain to alter CA-binding specificity. TRIMCyp from both rhesus and pigtailed macaques does not restrict HIV-1 infection and differs from human CypA at only two amino acid positions [D66N and R69H], both of which are outside the active site and differently affect binding to lentiviral capsids (50, 57). We previously demonstrated that CypA_D66N_ and CypA_D66N/R69H_ do not bind HIV-1 CA and that cells expressing these mutant CypA proteins phenocopy CsA-treated and CypA^-/-^ cells with regard to HIV-1 infection and MX2 sensitivity (44). We therefore engineered the corresponding changes of RANBP2 residues D3126 and K3129 (Fig. 3A) and tested these for their effect on HIV CA binding by generated cells expressing HA-tagged chimeric proteins in which the Cyp domain of owl monkey TRIMCyp was replaced with human RANBP2-Cyp (TRIM-BP2Cyp) containing D3126N, K3129H, or both D3126N/K3129H substitutions. TRIM-huCypA fusions with the corresponding D66N, R69H, and D66N/R69H substitutions (44) were included for comparison (Fig. 3B-C). We then challenged these cells with HIV-1_WT_, HIV-2, HIV-1 CA mutants, and other lentiviruses (Figs. 3B and *SI Appendix*, Fig. S4). HIV-1_WT_ infection was potently inhibited (∼100-fold) by TRIM-BP2Cyp and TRIM-BP2Cyp_K3129H_ and was not rescued by CsA addition as expected, while TRIM-BP2Cyp_D3126N_ and TRIM-BP2Cyp_D3126N/K3129H_ were inactive against HIV-1_WT_. This restriction profile was similar to the corresponding mutations in TRIM-huCypA fusions, suggesting that the same residues are involved in HIV-1 CA binding by CypA and RANBP2-Cyp.

To generate RANBP2 point mutant cell lines, we modified our approach for generating RANBP2_ΔCyp_ cells (*SI Appendix*, Fig. S1B) to express full-length RANBP2 with D3216N and/or K3129H mutations (or WT sequence as a control), followed by a P2A ribosomal skipping sequence and a hygromycin B-resistance gene (Hygro^R^). Single cell clones with the desired biallelic mutations were confirmed by sequencing of both genomic DNA and mRNA. As above, RANBP2-Cyp mutant cell lines were then transduced with a lentiviral vector for doxycycline-inducible expression of MX2 and infected with GFP-reporter viruses (Figs. 3D-G and *SI Appendix*, Fig. S2 and S3). Similar to the phenotype in RANBP2_ΔCyp_ cells, we observed a ∼2-fold reduction in infection and a reduction in MX2 sensitivity of HIV-1_WT_ in both RANBP2_D3126N_ and RANBP2_D3126N/K3129H_ cells. This effect was specific to mutations that affected binding by RANBP2-Cyp, since both infectivity and MX2 sensitivity of HIV-1_WT_ were similar in control and RANBP2_K3129H_ cells, and suggests that the effects of deletion of the Cyp domain of RANBP2 on HIV-1 infection and MX2 sensitivity are the direct result of abrogating CA-RANBP2-Cyp interactions (although we cannot definitively exclude the possibility that RANBP2_D3126N_ affects interactions with a cellular substrate relevant for HIV-1 CA-MX2 interactions). HIV-1_G89V CA_ was insensitive to all TRIM-Cyp fusions as expected (*SI Appendix*, Fig. S4), and the ability of MX2 to enhance HIV-1_G89V CA_ infection was unaffected in any of the RANBP2 point mutant cell lines (Figs. 3E), further indicating that the reduction in MX2 enhancement of HIV-1_G89V CA_ infection in RANBP2_ΔCyp_ cells (Figs. 2B) is the result of CA-independent effects.

The HIV-1 CA mutants N57A, N74D, and T210K exhibited similar sensitivity to the chimeric TRIM-Cyps as the WT virus (*SI Appendix*, Fig. S4). Accordingly, these CA mutants exhibited similar infectivity and MX2 sensitivity in RANBP2_ΔCyp_ and RANBP2-CA binding point mutant cell lines (*SI Appendix,* Fig S2A, B, and E). Notably, HIV-1_A92E CA_ and HIV-1_G94D CA_, which are CsA-dependent in HeLa cells, exhibited a distinct profile of sensitivity to TRIM-BP2Cyp vs. TRIM-Cyp fusions (*SI Appendix*, Fig. S4). Additionally, unlike HIV-1_WT_ and other CA mutants, infectivity and MX2 sensitivity of HIV-1_A92E CA_ and HIV-1_G94D CA_ in RANBP2 point mutant cell lines did not appear to correlate with the ability to bind RANBP2-Cyp (*SI Appendix,* Fig S2C-D), indicating that interactions between these CA mutants and CypA/RANBP2-Cyp are highly complex.

### Requirements for RANBP2-Cyp interactions are virus-specific

The D66N and R69H mutations in rhesus and pigtailed macaque TRIM-Cyp, which eliminate HIV-1 binding, increase affinity for HIV-2 CA (50), and HIV-2 therefore has a different sensitivity profile to TRIM-huCypA fusions compared to HIV-1 [Fig. 3B and (44)]. However, HIV-2 exhibited a similar profile as HIV-1 to sensitivity to TRIM-BP2Cyp fusions (Fig. 3B). Similarly, the sensitivity of SIVagmTAN to TRIM-BP2Cyp fusions with D3126N and/or K3129H mutations does not correspond with its sensitivity to TRIM-huCypA fusions with D66N and/or R69H mutations (*SI Appendix*, Fig. S4). These results indicate that, in contrast to HIV-1, binding to HIV-2 and SIVagmTAN CAs by CypA and RANBP2-Cyp involve distinct residues.

While mutation of D3126N abrogated the sensitivity of HIV-2 to TRIM-BP2Cyp restriction, the infectivity of HIV-2 was only slightly reduced in RANBP2_D3126N_ and RANBP2_D3126N/K3129H_ cells (Fig. 3F), suggesting that other residues are responsible for HIV-2 interactions with RANBP2-Cyp. HIV-2 also remained sensitive to MX2 activity in all RANBP2 mutant cells, similar to both SIVmac and SIVagmTAN (Fig. 3F and *SI Appendix*, Fig. S3A-B). These findings indicate that while restriction of HIV-1 by MX2 requires both CA-RANBP2-Cyp and CA-CypA interactions, restriction of HIV-2, SIVmac, and SIVagmTAN is independent of CA-Cyp interactions. Additionally, as expected, EIAV and FIV infection and MX2 sensitivity were largely unaffected in RANBP2 mutant cells (*SI Appendix*, Fig. S3C-D). Overall, these experiments indicate that direct interactions between RANBP2-Cyp and the viral CA affect infectivity and MX2 sensitivity in a CA sequence-dependent manner.

### HIV-1 integration is mislocalized in RANBP2ΔCyp cells

HIV-1 integration, which favors active transcription units within gene-dense regions and speckle-associated domains (SPADs), and disfavors heterochromatin regions such as lamina-associated domains (LADs), is mediated by interactions between HIV-1 integrase and CA with multiple host cell proteins [reviewed in (58, 59)]. Previous reports have indicated opposing effects of RANBP2 and CypA on HIV-1 integration in HeLa cells, with RANBP2 knockdown leading to reduced integration in gene dense regions and CpG islands (23) and CsA treatment (or infection with G89V/P90A CA mutants) leading to increased integration in gene dense regions, CpG islands, and SPADs, and a decrease in integration in LADs (5, 60). MX2 has also been shown to affect integration patterns in HOS cells, slightly reducing genic integration, and integration surrounding transcriptional start sites (TSSs) and CpG islands (33). To determine how the RANBP2-Cyp domain specifically affects HIV-1 integration targeting, we assessed integration in CypA-deficient (44), RANBP2_ΔCyp_, and control cells in the presence or absence of MX2 expression (Fig. 4 and *SI Appendix,* Dataset S1). Cells were infected with WT HIV-1 containing a heterologous sequence embedded in the U3 region to enable discrimination from the lentiviral vector for doxycycline-inducible MX2 expression (61). Proviral integration sites were amplified, sequenced, and analyzed for association with genes, gene density, SPADs, LADs, TSSs, and CpG islands as previously described (62, 63). We found that integration into genes was increased in both CypA^-/-^ and RANBP2_ΔCyp_ cells (Fig. 4A) as was integration into gene dense regions (>20 genes/Mb) (Fig. 4B), SPADs (Fig. 4D), and CpG islands (Fig. 3F), while there were fewer integration sites in LAD-associated DNA (Fig. 4E) and no change in integration within TSSs (Fig. 4C). MX2 expression resulted in reduced targeting to gene dense regions and SPADs, with a concomitant increase in integration in LADs (although the effects were not statistically significant for gene density/SPADs in control cells, there was a clear trend). Notably, although the ability of MX2 to restrict HIV-1 infection is reduced in CypA^-/-^ and RANBP2_ΔCyp_ cells [(44) and Fig. 2], we found that the effect of MX2 on integration targeting was maintained in the absence of CA-Cyp interactions (Figs. 4B, D, E). These findings suggest that while CA-Cyp interactions are required for MX2-mediated restriction, MX2 also affects the interactions of HIV-1 CA with other cellular factors involved in integration targeting (e.g., Nups, CPSF6) and/or the kinetics of nuclear import independently of CA-Cyp interactions.

**Fig. 4.**
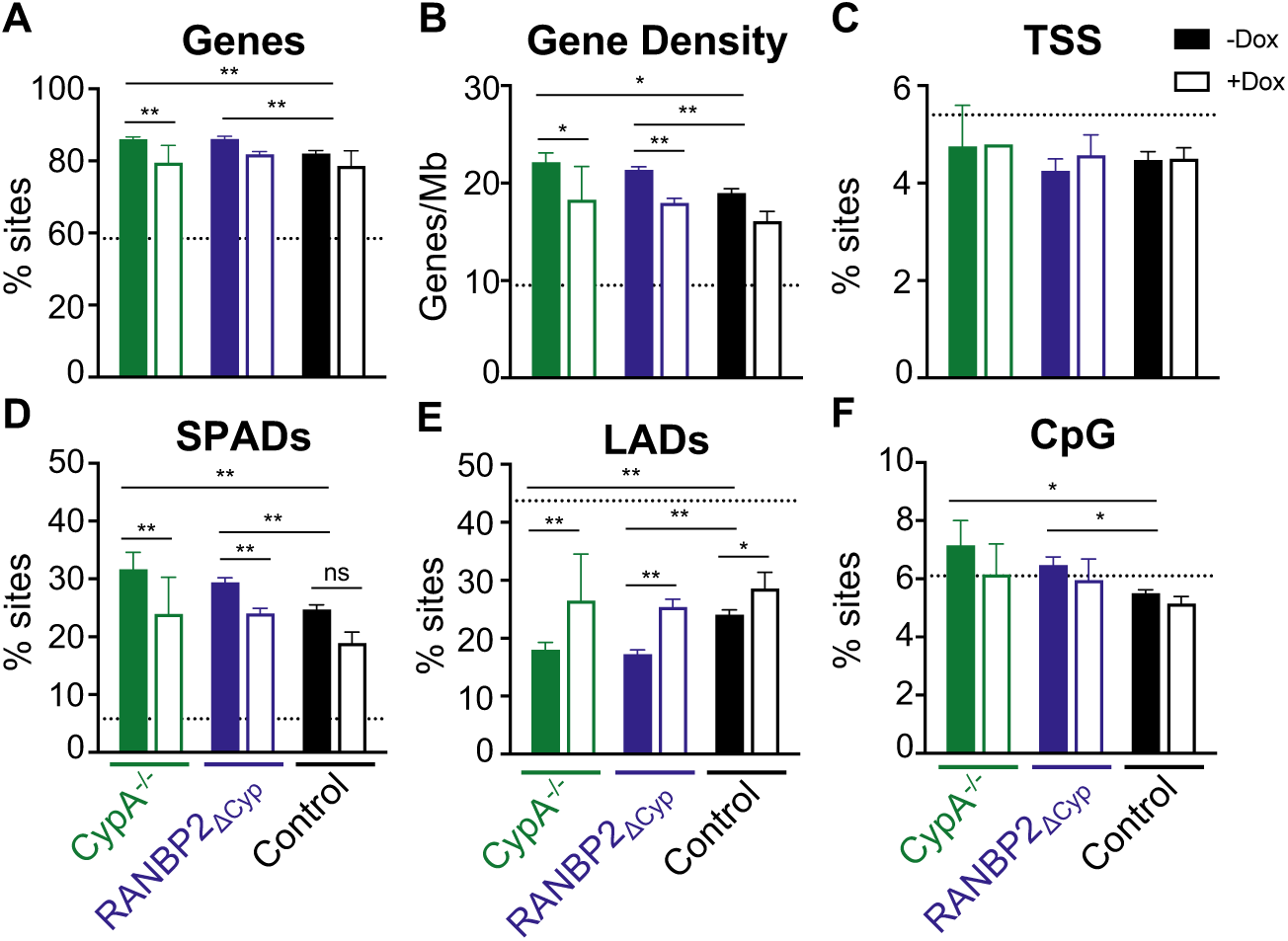
Similar effects of CypA knockout and RANBP2-Cyp deletion on HIV-1 integration site targeting. HIV-1 integration sites in unmutated control, CypA^-/-^, and RANBP2_ΔCyp_ HT1080s stably transduced with doxycycline-inducible MX2 in the presence (open bars) or absence (filled bars) of doxycycline were analyzed for (*A*) genes, (*B*) gene density, (*C*) transcription start sites (TSSs), (*D*) speckle-associated domains (SPADs), (*E*) Lamina-associated domains (LADs), and (*F*) CpG islands. Dashed lines indicate random integration control values. Infection and integration analysis was performed in duplicate in one representative CypA^-/-^ and control, and two RANBP2_ΔCyp_ clones. Control and RANBP2_ΔCyp_ cells were analyzed in two independent experiments. Comparisons between infection conditions were analyzed by Fisher’s exact test (*A, C-F*) or Wilcoxon rank-sum test (*B*). (*P*≥0.05); * *P* < 0.05; ** *P* < 0.0001.

### Deletion of the RANBP2-Cyp domain alters nucleoporin requirements for HIV-1 infection and MX2 sensitivity

Our previous work demonstrated that multiple Nups functionally interact in complex ways that affect the efficiency of HIV infection and the antiviral activity of MX2 (18). Therefore, we next investigated whether the RANBP2-Cyp domain affects the requirements for other nucleoporins using our previously validated siRNA library targeting Nups and NTRs [Fig. 5A and (18)]. Despite the previously discussed complications of Nup depletion, this approach remains valuable for the identification of changes in viral dependence upon Nups under various conditions (eg. comparison of WT versus CA mutant viruses or WT versus mutant cells) and can provide valuable insights for more in-depth future investigation of individual Nups. Control and RANBP2_ΔCyp_ cells were transfected with siRNA before being split into replicate wells for MX2 induction (via doxycycline addition) followed by infection with HIV-1_WT_ (Fig. 5B-C and *SI Appendix*, Figs. 5 and 6A-B). Deletion of the RANBP2-Cyp domain did not result in global changes to HIV-1_WT_ sensitivity to nucleoporin depletion, as most Nup and NTR depletions had similar effects on HIV-1_WT_ infection in both RANBP2_WT_ and RANBP2_ΔCyp_ cells (Fig. 5B). There were, however, a number of differences in Nup requirements for HIV-1_WT_ infection in RANBP2_ΔCyp_ cells, including a dramatic increase in sensitivity to NUP155 knockdown, and decreased sensitivity to NUP107, NUP93, and RANBP2 depletion. Additionally, while Nup62 subcomplex depletion modestly enhanced HIV-1_WT_ infection in control cells, depletion of members of this subcomplex slightly reduced infectivity in RANBP2_ΔCyp_ cells. Many Nup/NTR depletions also had similar effects on MX2 activity against HIV-1_WT_ in both control and RANBP2_ΔCyp_ cells (Fig. 5C) (e.g., increased upon KPNB1 depletion, decreased upon SEH1, NUP107, NUP133, ELYS, and TNPO1 depletion). However, there were many Nup depletions that altered the sensitivity of HIV-1_WT_ to MX2 in RANBP2_ΔCyp_ cells, most notably a total loss in MX2 sensitivity upon NUP155 depletion in RANBP2_ΔCyp_ cells, which enhanced antiviral activity in wild-type cells.

**Fig. 5.**
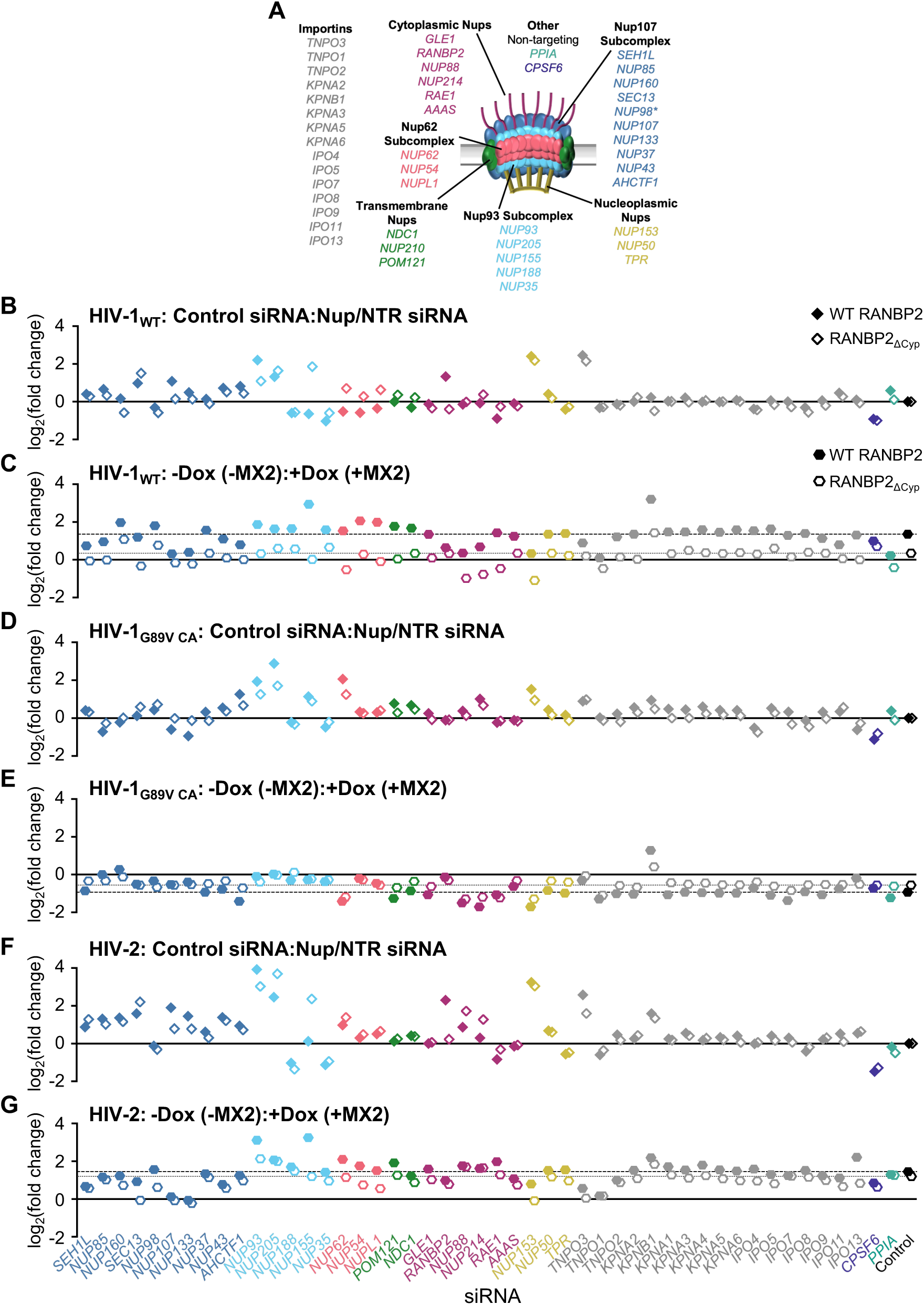
Effect of RANBP2-Cyp on the nucleoporin requirements for HIV infection and MX2 sensitivity. (*A*) Schematic representation of the nuclear pore complex and genes included in siRNA library color coded by subcomplex. (*- NUP98 is listed as a member of Nup107 subcomplex, however. Nup98 and Nup96 are produced following autoproteolytic cleavage of a polyprotein precursor (80, 81), this siRNA will target both Nups). Importins/nuclear transport receptors (NTRs) included in the siRNA library listed in grey. Also included, siRNA targeting CypA, CPSF6, and a non-targeting control siRNA. (*B, D, F*) Effect of Nup/NTR depletion on (*B*) HIV-1_WT_, (*D*) HIV-1_G89V CA_, and (*F*) HIV-2 infection in control (filled diamonds) and RANBP2_ΔCyp_ (open diamonds) cells shown as a ratio [log_2_(fold change of control siRNA/Nup/NTR siRNA)]. Average fold change calculated from eight technical replicates from two independent experiments. (*C, E, G*) Effect of Nup/NTR depletion on MX2 sensitivity of (*C*) HIV-1_WT_, (*E*) HIV-1_G89V CA_, and (*G*) HIV-2 in control (filled hexagons) and RANBP2_ΔCyp_ (open hexagons) cells shown as a ratio [log_2_(fold change of -Dox(-MX2)/+Dox(+MX2))]. Dashed line represents effect in control cells dotted line represents effect in RANBP2_ΔCyp_ cells; siRNAs are color coded by subcomplex as in *A*.

As expected, sensitivity of HIV-1_G89V CA_ to nucleoporin depletion was also largely unaffected by Nup depletion (Fig. 5D and *SI Appendix* Figs. 6C-D), although we did observe some changes in sensitivity to NUP107, NUP133, NUP93, NUP205, and NUP62 depletion, indicating that the Cyp domain may affect the function/conformation of other Nups. Similarly, MX2-mediated enhancement of HIV-1_G89V CA_ infectivity was amplified and diminished by similar Nup depletions in both control and RANBP2_ΔCyp_ cells (Fig. 5E) (e.g., amplified by NUP62, NUP88, NUP214, RAE1, and NUP153 depletion and diminished by NUP85, NUP160, Nup93 subcomplex, NUP54, RANBP2, and TNPO3 depletion). Notably, while the antiviral activity of MX2 against HIV-1_WT_ was reduced in control cells upon depletion of the cytoplasmic ring Nups, NUP88, NUP214, and RAE1 and upon depletion of NUP153, MX2 expression enhanced HIV-1_WT_ infection upon these knockdowns in RANBP2_ΔCyp_ cells, similar to its effect on HIV-1_G89V CA_ in both control and RANBP2_ΔCyp_ cells (Fig. 5B-E). MX2 also modestly enhanced HIV-1_WT_ infection upon NUP62, RAE1, and TNPO1 knockdown. These results suggest that the amplification of MX2-dependent increases in HIV-1_G89V CA_ infectivity upon several Nup depletions results from a failure of this mutant to interact with RANBP2-Cyp and may therefore be independent from the effects of CypA on nuclear import and MX2 sensitivity.

Finally, we investigated sensitivity of HIV-2 to Nup/NTR depletion in RANBP2_ΔCyp_ cells (Fig. 5F-G and *SI Appendix,* Fig S6E-F). As was the case for HIV-1, most knockdowns had similar effects on HIV-2 infection in control and RANBP2_ΔCyp_ cells. Like HIV-1, sensitivity of HIV-2 to NUP155 depletion was dramatically increased in RANBP2_ΔCyp_ cells, as was elimination of sensitivity to RANBP2 depletion (Fig. 5B and F). We also observed some HIV-2-specific effects of Nup/NTR depletion (e.g., increased sensitivity to NUP205, NUP88, and NUP214 knockdown and decreased sensitivity to TNPO3 knockdown). HIV-2 sensitivity to MX2 was retained in RANBP2_ΔCyp_ cells (Fig. 2A-B), and while some Nup depletions reduced the susceptibility of HIV-2 to MX2 in RANBP2_ΔCyp_ cells (e.g., SEC13, NUP98, NUP93, NUP155, the Nup62 subcomplex, and NUP153) (Fig. 5G), we did not observe any instances in which MX2 enhanced HIV-2 infectivity of RANBP2_ΔCyp_ cells, as it did for HIV-1. Collectively, these findings indicate that deletion of the RanBP2-Cyp domain alters the nuclear import pathways available to HIV (or nuclear import kinetics) predominantly via its effects on CA-specific interactions with other NPC components; and further suggests that alterations in MX2 sensitivity are determined by these Cyp-domain-dependent changes in nuclear import pathway usage, kinetics, and/or CA conformation.

## Discussion

Interactions with numerous host proteins are required for retroviral access to the nucleus and for integration. Nups appear to play a key role in both HIV-1 passage across the nuclear envelope, and in downstream interactions that affect viral access to the host chromatin. Complexities in NPC composition have made determining the functionally relevant interactions between HIV-1 CA and Nups a significant challenge. This difficulty extends to our understanding of MX2 activity, which is clearly affected by complex interactions with its cellular environment. Our previous work that employed a systematic knockdown approach indicated that multiple Nups are involved in both HIV-1 infection and the antiviral activity of MX2 (18). However, data obtained using knockdown/knockout approaches must be interpreted with caution due to the existing variability in NPC composition, and the significant pleiotropic effects of many Nup depletions (18). Similarly, studies establishing direct interactions between Nups and CA either by demonstrating susceptibility to chimeric TRIM5-fusion proteins, or by *in vitro* biochemical approaches, may or may not indicate that such interactions have functional consequences in the context of virus infection. Modern gene-editing technology now provides the opportunity to directly test the role of individual Nup domains in HIV-1 infection and MX2 activity by generating cells expressing Nups from their endogenous loci, but with specific domains deleted. This precise construction of deletion mutants allows for the modification of the functional capabilities of the NPC without disrupting its assembly or altering overall expression levels. Using this approach, we investigated the function of the Cyp homology domain of the cytoplasmic filament Nup RANBP2 in HIV-1 infection and MX2 activity. We demonstrate that HIV-1 CA-Cyp domain interactions are not required for infection but affect integration site targeting and MX2 activity in a manner similar to CA-CypA interactions (Figs. 2A-B, 3D-E, and 4). We also determined that the RANBP2-Cyp domain affects the requirements for other Nups for infection and MX2 sensitivity (Fig. 5).

The antiviral activity of MX2 against HIV-1 is reduced in cells containing mutations in CypA that affect CA recognition (44), and in this report, we demonstrate that similar mutations in RANBP2-Cyp reduce the antiviral activity of MX2. Importantly, however, the effect of MX2 on integration site targeting was retained in both CypA^-/-^ and RANBP2_ΔCyp_ cells (Fig. 4). A recent preprint proposed that MX2 restricts HIV-1 (and herpes simplex virus-1) by forming biomolecular condensates that act as a nuclear pore decoy (64); the authors further suggest that MX2-resistance of the P90A CA mutant is the result of its lower propensity to accumulate within these condensates. Our findings reported here would indicate that HIV-1 CA interacts with MX2 regardless of its conformation/interaction with other cellular cofactors, whether this interaction results in abortive infection or altered integration targeting is determined (at least in part) by CA-Cyp interactions. This is also consistent with our previous work demonstrating that MX2-resistance is context dependent and is affected by CA mutations, cell-type, cell-cycle, and CypA (18, 44) and that MX2 can increase infection of HIV-1 in some contexts [Fig. 5 and (18)].

Knockdown of RANBP2 reduces HIV-1 integration into gene dense regions, similar to cells lacking CPSF6 (or CA mutants that do not bind CPSF6) (23, 63, 65, 66), however, we demonstrate here that this effect is not mediated by CA-RANBP2-Cyp interactions, as integration is mislocalized in RANBP2_ΔCyp_ cells in a manner similar to CypA^-/-^ cells (increased targeting to gene dense regions and increased SPAD targeting) (Fig. 4). Given the increasing evidence that CypA is stripped from the capsid at the nuclear pore (67, 68), and that CypA-CA interactions affect nuclear entry and integration in a CPSF6-dependent manner (60), one possibility is that sequential binding of CA to CypA, followed by RANBP2-Cyp, affects downstream interactions with CPSF6. While CPSF6 does not appear to play a major role in MX2 activity (Fig. 5 and (18)), it has been proposed that CPSF6 and MX2 cooperate to prevent interactions with RANBP2 (69). Deletion of RANBP2-Cyp abrogated the sensitivity of HIV-1_WT_ and HIV-2 to RANBP2 knockdown, similar to the phenotype observed for Cyp-binding mutant CAs in cells expressing WT RANBP2 [(30, 32, 34–36) and Fig. 5). Interestingly, CsA treatment also rescues HIV-1 infection from RANBP2 (and NUP153) knockdown in some cell types (70), suggesting that CA-interactions with both CypA and the RANBP2-Cyp domain determine the requirement for RANBP2. Similarly, RANBP2-Cyp-CA interactions appear to determine the requirement for NUP155, as HIV-1_G89V CA_ is sensitive to NUP155 depletion, and both HIV-1_WT_ and HIV-2 become highly sensitive to NUP155 knockdown in RANBP2_ΔCyp_ cells (Fig. 5). NUP155 is an inner ring Nup that lacks an FG domain (71) and does not bind HIV-1 CA tubes *in vitro* (18). It will therefore be of interest in future investigations to determine how NUP155 affects HIV-1 and MX2 without directly interacting with CA [perhaps by affecting the structural conformation of individual nuclear pores (72, 73)].

In summary, this report demonstrates a specific role for the Cyp domain of RANBP2 in HIV infection and MX2 sensitivity, and further indicates that precise manipulation of NPCs is key to revealing the functional interactions that determine the outcome of HIV-1 infection. Furthermore, this approach should be of great value in revealing the details of normal cellular nucleocytoplasmic trafficking.

## Materials and Methods

### Cell lines and viruses

Culture of cell lines has been previously described (18, 44). Generation of viral stocks and infectivity experiments have been previously described (18, 44, 61, 74) (for details, see *SI Appendix, Supplementary Methods*).

### Generation of RANBP2_ΔCyp_ and point mutant cell lines

RANBP2 mutant cell lines were generated via CRISPR/Cas9-mediated genome editing using AAVs for delivery of donor sequences for homology directed repair as previously described (44). Details are provided in the *SI Appendix, Supplementary Methods*.

### CA-binding assay with HIV-1 CA tubes

Binding assays with HIV-1 CA nanotubes were performed as previously described (75–77), details are provided in the *SI Appendix, Supplementary Methods*.

### RNA interference

RNA interference targeting Nups and NTRs was performed as previously described (18), (for details, see *SI Appendix, Supplementary Methods*).

### Western blotting and immunofluorescence

Detection of proteins via western blotting and immunofluorescence were performed as previously described (18, 44), for details see *SI Appendix, Supplementary Methods*.

### Integration site analysis

Genomic DNA from HIV-1 infected cells was used to prepare libraries for integration site analysis as previously described (33, 78). Sequenced products were analyzed for genomic features and coordinates of integrated proviruses were identified as previously described (61, 63). For details, refer to the *SI Appendix, Supplementary Methods*. Illumina FASTQ raw reads are available at the National Center for Biotechnology Sequences Read Archive (Accession Number: BioProject PRJNA1147214)

### Statistical analyses

Statistical significance for CA binding and infectivity assays was determined using GraphPad software (two-way ANOVA or t-tests where appropriate). Comparison of percent integration was analyzed by Fisher’s Exact test; comparison of gene density was analyzed by Wilcoxon rank-sum test. Statistical analyses not included in Figures are included in *SI Appendix, Dataset S2*.

## Acknowledgements

We thank Drs. Fabian Schmidt and Yiska Weisblum consultation regarding design and production of AAV templates for homology-directed recombination, Robert Z. Zhang for helpful discussion, and Mariah Cashbaugh for technical support. Color-blind safe “muted” qualitative color schemes from Paul Tol were used for Figs. (79). This work was supported by grants from the NIH: R01AI1621172 and R01AI150988 (to M.Kane), R01 AI162665 (to M.K.), R01 AI052014 (to A.N.E.), and U54AI170791 (to M.Kane and A.N.E.). The content is solely the responsibility of the authors and does not necessarily represent the official views of the National Institutes of Health.

## Notes

### Competing Interest Statement

The authors have declared no competing interest.

