## Supporting Information for "Interplay between the cyclophilin homology domain of RANBP2 and MX2 regulates HIV-1 capsid dependencies on nucleoporins"

**This PDF file includes:**

Supplementary Methods
Figures S1 to S6
Tables S1 to S3
SI References

**Other supporting materials for this manuscript include the following:**

Dataset S1
Dataset S2

### Supporting Information Text
### Supplementary Methods

#### Plasmid construction

*TRIM5-fusions*
An LHCX MLV vector (Clontech) was engineered to express the N-terminal domains (RING, B-Box and coiled coil) from owl monkey (omk) TRIMCyp followed by an HA tag and cloning sites (*NotI*/*SalI*) that allowed introduction of various proteins as described in (1, 2). TRIM-huCypA fusions were previously reported (3). Plasmids expressing TRIM5-HA N-terminus fused to the human RANBP2-Cyp domain, or the human RANBP2-Cyp domain with the D3126N, K3129H, or D3126N/K31299H changes, were generated by overlap PCR with the primers indicated in Table S1.

*AAV donors*
To construct the pAAV-*RANBP2* donor plasmid for the generation of RANBP2$_{\Delta Cyp}$ cells (Fig. S1A), a 1965 bp region of the *RANBP2* locus lacking the Cyp domain and intron 28 (bases 62242 to 63214 and 64454 to 65446 NCBI Reference Sequence: NG_012210.2) was synthesized as a gBlock fragment (IDT) and inserted into the pAAV packaging plasmid (Cell Bio Labs) using *Nhe*I and *Xho*I. The PAM site for crRNA 1 was mutated from TGG to TGA and for crRNA to from GGG to GCC. For pAAV-*RANBP2* donor plasmids for RANBP2 point mutant cell lines, a gBlock fragment was synthesized containing an 890bp region of the *RANBP2* locus lacking intron 28 (bases 62831 to 63417 and 64151 to 64453) followed by a P2A cleavage site and Hygromycin-resistance gene, and 57 bases of exon 29 of *RANBP2* following the stop codon (bases 64454 to 64510) was inserted into pAAV-RANBP2$_{\Delta Cyp}$ using *BamHI* and *HindIII*. A silent *KpnI* site was introduced into intron 27 as shown in Fig. S1B. The wild-type donor template was synthesized, then RANBP2$_{D3126N}$, RANBP2$_{K3129H}$, and RANBP2$_{D3126N/K3129H}$ donors were generated via overlap PCR with the primers indicated in S1 table and inserted into pAAV-RANBP2-2A-Hygro using *KpnI* and *NdeI*.

#### Cell lines
HEK 293T, HeLa, and HT1080 cell lines were maintained in Dulbecco's Modified Eagles Medium [(DMEM) Gibco] with 10% fetal calf serum (Gibco), gentamicin (Gibco), and Plasmocin prophlyactic (InvivoGen). Plasmocin was not present in cultures during infection, transduction, or transfection. Cells were purchased from ATCC and were assumed to be authenticated by their supplier and were not further characterized. Cells were monitored quarterly for retroviral contamination by SYBR-Green based PCR RT assay (4, 5) and tested for mycoplasma contamination by MycoStrip detection kit (InvivoGen). Derivatives of HeLa and HT1080 cells containing doxycycline-inducible MX2 or fusion proteins were generated by transduction with LKO-derived lentiviral vectors (6) followed by selection in 1 µg ml$^{-1}$ puromycin (Sigma-Aldrich). HeLa cells stably expressing TRIM fusions were generated by transduction with LHCX-derived vectors followed by selection in 100 µg ml$^{-1}$ hygromycin B (Gibco). Vector stocks for transduction were generated by co-transfection of 293T cells with a VSV-G expression plasmid, an HIV-1$_{NL4-3}$ Gag-Pol expression plasmid, and LKO-derived vector, or an MLV Gag-Pol expression plasmid and an LHCX-derived vector using polyethyleneimine (PolySciences). Expression was induced in pLKO transduced cell lines through an overnight treatment with 500 ng/ml doxycycline hyclate (Sigma-Aldrich) prior to challenge with retroviruses or retroviral vectors.

#### Viruses
All viruses were generated by transfection of 293T cells using polyethyleneimine (PolySciences). GFP reporter proviral plasmids HIV-1$_{NL4-3}$ΔEnv-GFP (HIV-1, HIV-1$_{G89V}$, HIV-1$_{A92E}$, and G94D CA mutants (7)), NHGCapNLNM (wild-type, HIV-1$_{N57S}$, HIV-1$_{N57A}$, HIV-1$_{N57D}$, HIV-1$_{N74D}$, and HIV-1$_{T210K}$ CA mutants (8, 9), HIV-2$_{ROD}$ΔEnv-GFP, SIV$_{MAC}$ΔEnv-GFP, SIV$_{AGM}$TANΔEnv-GFP (10), and HIV-1$_{NL4-3}$ΔEnv-Luc-U3tag (11) (10 µg) were co-transfected with 1 µg of VSV-G expression plasmid. For EIAV and FIV, three plasmid vector systems (12, 13) were used to generate GFP reporter viruses, whereby 5 µg of Gag-Pol, 5 µg of packageable genome, and 1 µg of VSV-G expression plasmids were co-transfected.

#### Infection assays
Infectivity was measured in HeLa or HT1080 cells seeded in 96-well plates at 5 x 10$^3$ cells per well and inoculated with serial-dilutions of VSV-G pseudotyped GFP reporter viruses in the presence of 4 µg ml$^{-1}$

polybrene (Sigma-Aldrich). Where indicated, cyclosporine A (Sigma-Aldrich) was added to the cultures at the time of infection at 5 µM. Infected cells (%GFP positive of viable cells) were enumerated by FACS analysis using an Attune NxT coupled to an Autosampler (Invitrogen).

**Generation of RANBP2$_{\Delta Cyp}$ and point mutant cell lines**

AAVs containing donor sequence for homology directed repair were generated by co-transfection of 293T cells with pAAV-*RANBP2*, pAAV-RC (DJ), and pHelper (CellBio Labs) at a 1:1:1 ratio using polyethyleneimine (PolySciences). Supernatant was collected 72 h post-transfection. 1x$10^5$ HT1080 cells in a 24-well plate were infected with 200 µL of AAV-containing supernatant 4-6 h prior to reverse transfection with Cas9 ribonucleoprotein complexes (RNPs). Custom Alt-R$^{TM}$ CRISPR Cas9 guide RNAs targeting the *RANBP2* locus were designed using the IDT website. RNPs containing crRNA 1 or 2:tracerRNA duplexes were generated according to manufacturer instructions: 1 µL of 100 µM crRNA, 1 µl of 100 µM tracrRNA-ATTO 550 (IDT) in 98 µL of Nuclease-Free Duplex Buffer (IDT) were heated at 98$^o$C for 5 min, then cooled to room temperature. 15 µL of annealed guide RNA oligos were mixed with 15 µL of 1 µM recombinant Cas9 (IDT) in 220 µL of Opti-MEM media (Gibco) and incubated for 5 min at room temperature, complexes were used immediately or stored at -80$^o$C. HT1080 cells were reverse transfected in 96-well plates with RNPs using RNAiMax (Thermo Fisher) according to manufacturer instructions, transfection complexes containing 15 µL of each RNP was mixed with 1.2 µL of RNAiMax and 18.8 µL of OptiMEM for 20 min at room temperature, followed by addition of 4x$10^4$ cells/well. For generation of point mutant cell lines, 1 µM Alt-R$^{TM}$ HDR Enhancer V2 (IDT) was included. RANBP2$_{\Delta Cyp}$ control cells were transfected with RNPs containing the IDT negative control crRNA. 16 h post-transfection, cells were transferred to a 24-well plate, and AAV infection/transduction was repeated after an additional 24-48 h. Single-cell clones were then derived by limiting dilution. For point mutant and wild-type control cell lines, bulk populations were placed in Hygromycin selection for five days prior to seeding clones.

RANBP2$_{\Delta Cyp}$ clones were screened via PCR amplification from genomic DNA using BP2delCyp Check HR F and R primers (see Table S1). RANBP2 point mutant clones were screened via PCR amplification from genomic DNA using BP2 Exon 26 F1 and BP2 Cyp R followed by *KpnI* digestion. Candidates were confirmed by PCR and sequencing from both genomic DNA and cDNA [using SuperScript$^{TM}$ III Reverse Transcriptase (Thermo Fisher)]. Genomic DNA and mRNA were extracted using NucleoSpin Tissue and NucleoSpin RNA kits (Macherey-Nagel), respectively. Candidates were confirmed by western blot and sequencing from genomic DNA. Nine RANBP2$_{\Delta Cyp}$ and three control clones were selected for experiments; each experiment included three to four RANBP2$_{\Delta Cyp}$ and two to three control clones. Two homozygous RANBP2$_{WT\text{-}2A\text{-}Hygro}$, two RANBP2$_{D3126N}$, three RANBP2$_{D3126N/K3129H}$, and one RANBP2$_{K3126H}$ clones were identified and selected for experiments. Experiments show data from multiple clones combined, or one representative clone (as we previously reported, HT1080 cells exhibit limited clonal variability (3)).

**CA-binding assay with HIV-1 CA tubes**

CA(WT), CA(G89V), and CA(N57A) were expressed from pET3a in BL21-DE3 cells and purified as previously described (14-16). CA nanotubes were assembled in a high-ionic strength buffer (15 mM Tris-HCl, pH 8.0; 2 M NaCl) as described (17, 18). Binding assays with HIV-1 CA nanotubes were performed as described (9). In brief, control and RANBP2$_{\Delta Cyp}$ HT1080 cells were lysed by adding a passive lysis buffer (Promega) supplemented with protease inhibitor cocktail (Roche). NaCl concentration in lysates was adjusted to 2 M and lysates were centrifuged at 13,000 x g for 2 min at 4℃. Then the supernatant of the cell lysates was added to the preformed CA nanotubes and incubated at room temperature for 30 min. Following centrifugation at 13,000 x g for 2 min at 4℃, the supernatant was saved, and the pellet was washed three times with the high-ionic strength buffer. LDS Reducing Sample buffer (Thermo Scientific Chemicals) was added to both pulled-down and unbound fractions, and they were subjected to SDS-PAGE. The proteins of interest were detected by immunoblotting using respective antibodies.

**RNA interference**

RNA interference using a custom siRNA library (Table S2) targeting Nups and NTRs was performed as previously described (9). RANBP2$_{\Delta Cyp}$ or control HT1080 cells stably transduced with doxycycline-inducible MX2 or were reverse transfected with 25 pmol of siRNA (SMARTpool, Dharmacon; SI Appendix S2 Table) using Lipofectamine RNAiMax (Invitrogen) at a concentration of 5 x $10^4$ cells/ml in 12-well plates. Non-targeting siRNA were used as controls, no significant difference in viral infectivity or Mx2-restriction were observed for each of these controls, as such, for each experiment, only the non-targeting siRNA control is shown. 24 h after transfection, cells were trypsinized, diluted 1:2.5 and re-plated in 96-well plates and treated with doxycycline, followed by infection with GFP reporter viruses 36-48 h later (See Fig. S5A).

**Western blotting**

Cell suspensions were lysed in NuPage LDS sample buffer (Invitrogen), followed by sonication, and separated by electrophoresis on NuPage 4-12% Bis-Tris gels or 4-8% Tris-Acetate gels (Invitrogen) and blotted onto polyvinylidene fluoride (PDVF, BioRad Laboratories). Membranes were incubated with the antibodies listed in Table S3, followed by incubation with goat anti-rabbit-HRP or goat anti-mouse-HRP secondary antibodies (Jackson ImmunoResearch). SeeBlue Plus2 and HiMark Pre-stained Protein Standards (Thermo Fisher) were used. Blots were developed with SuperSignal West Femto Maximum Sensitivity Substrate (Thermo Fisher) and imaged on a C-Digit scanner (LI-COR Biosciences).

**Immunofluorescence**

RANBP2$_{\Delta Cyp}$ or control HT1080 cells stably transduced with doxycycline-inducible MX2-tagRFP were seeded onto an 8-chamber gelatin-coated glass coverslip (Ibidi) and treated with 500 ng/ml doxycycline 16 h prior to fixation with 4% paraformaldehyde. Cells were then permeabilized with 0.5%Triton X-100 (Thermo Scientific) and immunostained with the indicated antibodies (Table S3) followed by goat anti-mouse or goat anti-rabbit Alexa 488 or Alexa 647 secondary antibodies (Molecular Probes). DNA was stained with Hoescht 33342 (Thermo Scientific). Cells were visualized on an EVOS M7000 digital microscope (Thermo Scientific) at 100X magnification. Image generation and deconvolution analysis were completed with the Celleste software suite (Thermo Scientific).

**Integration site analysis**

CypA$^{-/-}$ (3), RANBP2$_{\Delta Cyp,}$ or control HT1080 cells stably transduced with doxycycline-inducible MX2 were infected with DNase-treated (Turbo DNase, ThermoFisher) U3-tagged HIV-1 at a multiplicity of infection (MOI) of one. Three days post-infection, genomic DNA was extracted using the Machery-Nagel NucleoSpin Tissue Kit. Integration libraries were prepared using ligation-mediated PCR (LM-PCR) essentially as previously described (19-21). Genomic DNA (5 µg) was digested overnight with a cocktail of enzymes (100 U each, *AvrII*, *NheI-HF*, *SpeI-HF*, and *BamHI-HF*) and purified using a PCR purification kit (GeneJET). Following overnight ligation (four parallel reactions) to asymmetric linkers, DNA was purified again using a PCR purification kit. The ligated samples were subjected to two rounds of LM-PCR using virus and linker-specific primers (Dataset S1). Following PCR purification, libraries were assessed for fragment size distribution by TapeStation-4150, quantified by DNA fluorimetry (Qubit), and then pooled at 10 nM. Pooled samples were further diluted to 2 nM in Illumina sequencing resuspension buffer (RSB, Ref#20762979). Next, PhiX Control v3 DNA was spiked at 30-40%, and the sample was diluted to 650 pM with RSB. The mixture (20 µl) was loaded into a P2 300 cycle cartridge and sequenced on an Illumina NextSeq 2000 sequencer.

Raw fastq files were demultiplexed using Sabre tool (22) or by a Perl script. Post demultiplexing, files were trimmed, aligned to human genome build hg19, and bed files were generated as described (11). Integration into genes and SPADs was scored as within these genomic coordinates. For TSSs, CpG islands, and LADs, sites were mapped within +/-2.5 kb windows (5 kb surrounding these coordinates). Gene density was assessed as number of genes per Mb. Random integration controls (RICs) were generated by shearing hg19 in silico using the restriction enzyme sites used to generate the wet-bench samples, and then mapping the resultant fragments with respect to the aforementioned genomic annotations (21).

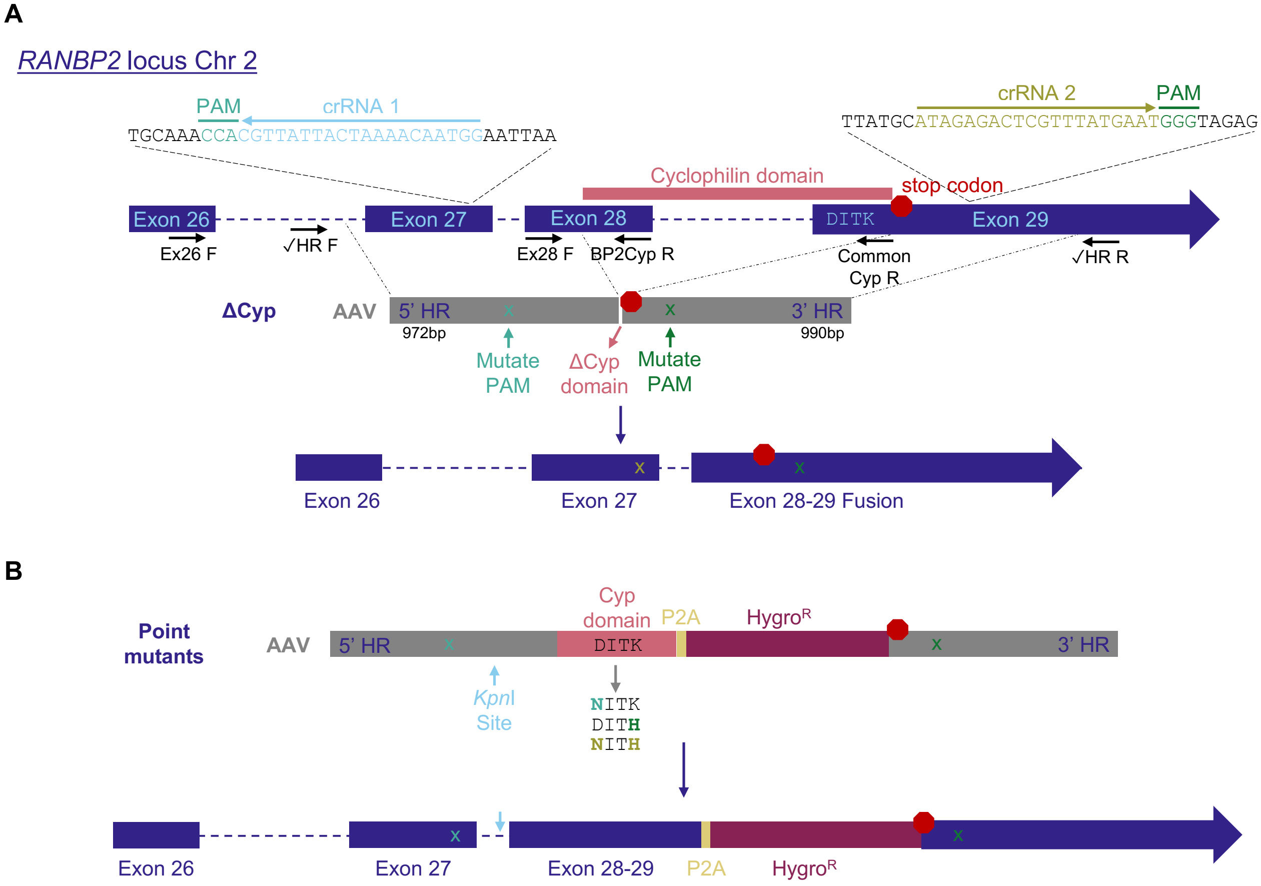


**Fig. S1. Generation of RANBP2 mutant cell lines**

A) Top - Schematic of the 3' end of the *RANBP2* locus on Chromosome 2 including location of the cyclophilin (Cyp) domain, residues 3126-3129, and stop codon indicated; crRNA guide targeting sites exon 27 and exon 29, and PCR primers for mutant clone screening and verification indicated. Diagram of AAV donor for homology-directed repair for generation of $RANBP2_{\Delta Cyp}$ cells shown below. Bottom – Schematic of the 3' of the locus following homology-directed repair with exons 28-29 fused and Cyp domain removed. B) Top - Diagram of AAV donor for generation of RANBP2 point mutant and control cells with silent *KpnI* site and amino acid residues in wild-type and mutant RANBP2 cells indicated. Bottom - Schematic of the 3' of the locus following homology-directed repair with exons 28-29 fused followed by P2A skipping site and Hygromycin-resistance gene ($Hygro^R$), the *KpnI* site in intron 27 indicated with a blue arrow.

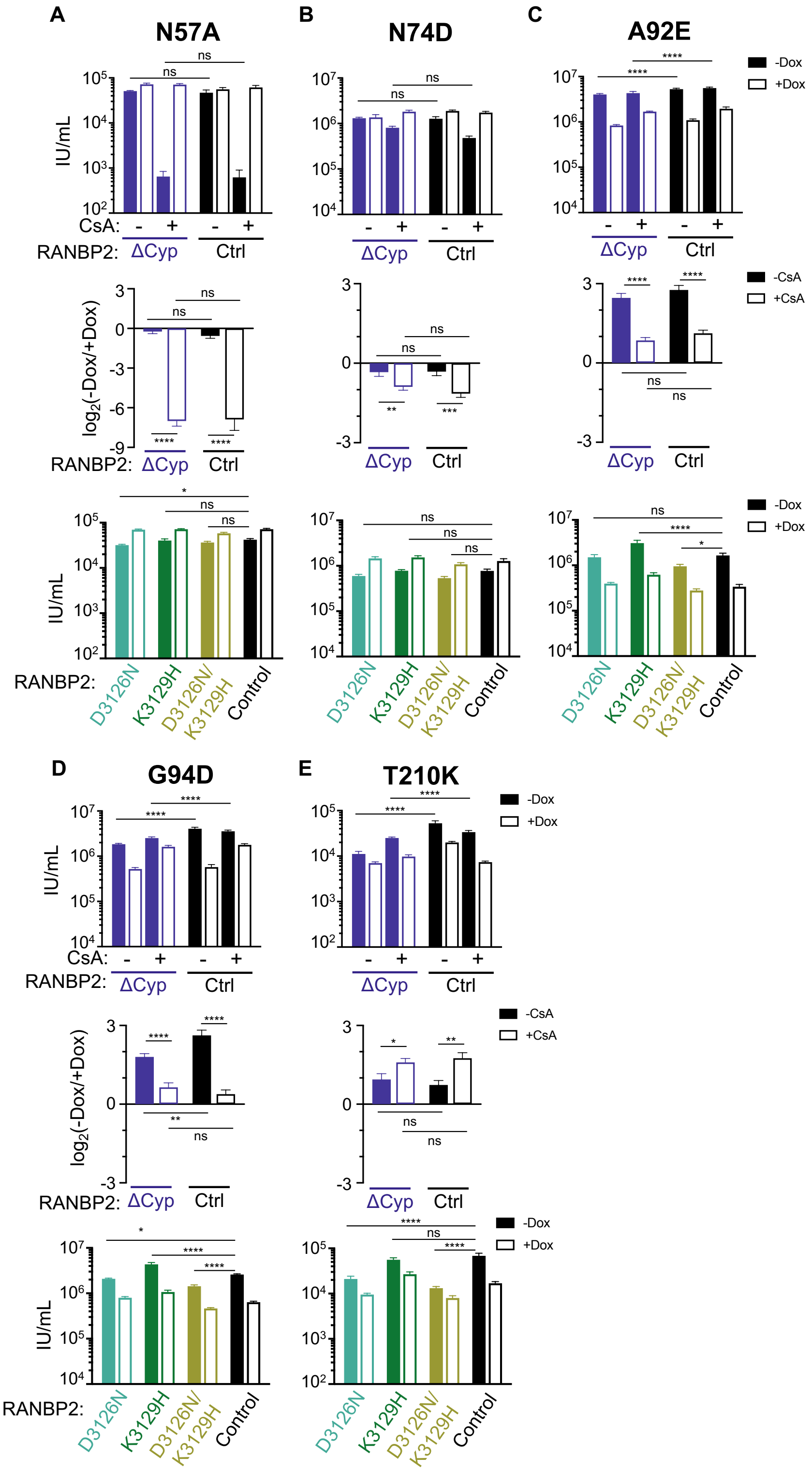


**Fig. S2. Effects of the RanBP2-Cyp domain on HIV-1 CA mutant infection and Mx2 sensitivity.**

A-E) (Top) Infection of control and RANBP2$_{\Delta Cyp}$ HT1080 cell clones stably transduced with doxycycline-inducible MX2 the presence (open bars) or absence (filled bars) of doxycycline and presence or absence of CsA with the indicated GFP reporter viruses. Titers are represented as mean + sem of infectious units (IU) per mL. RANBP2$_{\Delta Cyp}$: n≥12 technical replicates combined from ≥3 different clones; control: n≥8 technical replicates combined from ≥2 different clones; representative of ≥4 independent experiments. Statistical significance was determined by two-way ANOVA (Śídák's multiple comparisons test).

(Middle) Data from (top) shown as a ratio [fold change of -Dox(-MX2)/+Dox(+MX2)] in the presence (open bars) or absence (filled bars) of CsA. Average fold change calculated from three-four technical replicates per experiment; shown is mean + sem of $\log_2$(fold change) from four-eight independent experiments. Statistical significance was determined by paired *t* test.

(Bottom) Infection of control and RANBP2 point mutant HT1080 cell clones stably transduced with doxycycline-inducible MX2 the presence (open bars) or absence (filled bars) of doxycycline with the indicated GFP reporter viruses. Titers are represented as mean + sem of infectious units (IU) per mL. n≥12 technical replicates combined from ≥4 independent experiments. Statistical significance was determined by two-way ANOVA (Śídák's multiple comparisons test)

ns, not significant ($P \geq 0.05$); * $P < 0.05$; ** $P < 0.01$; *** $P < 0.001$; **** $P < 0.0001$.

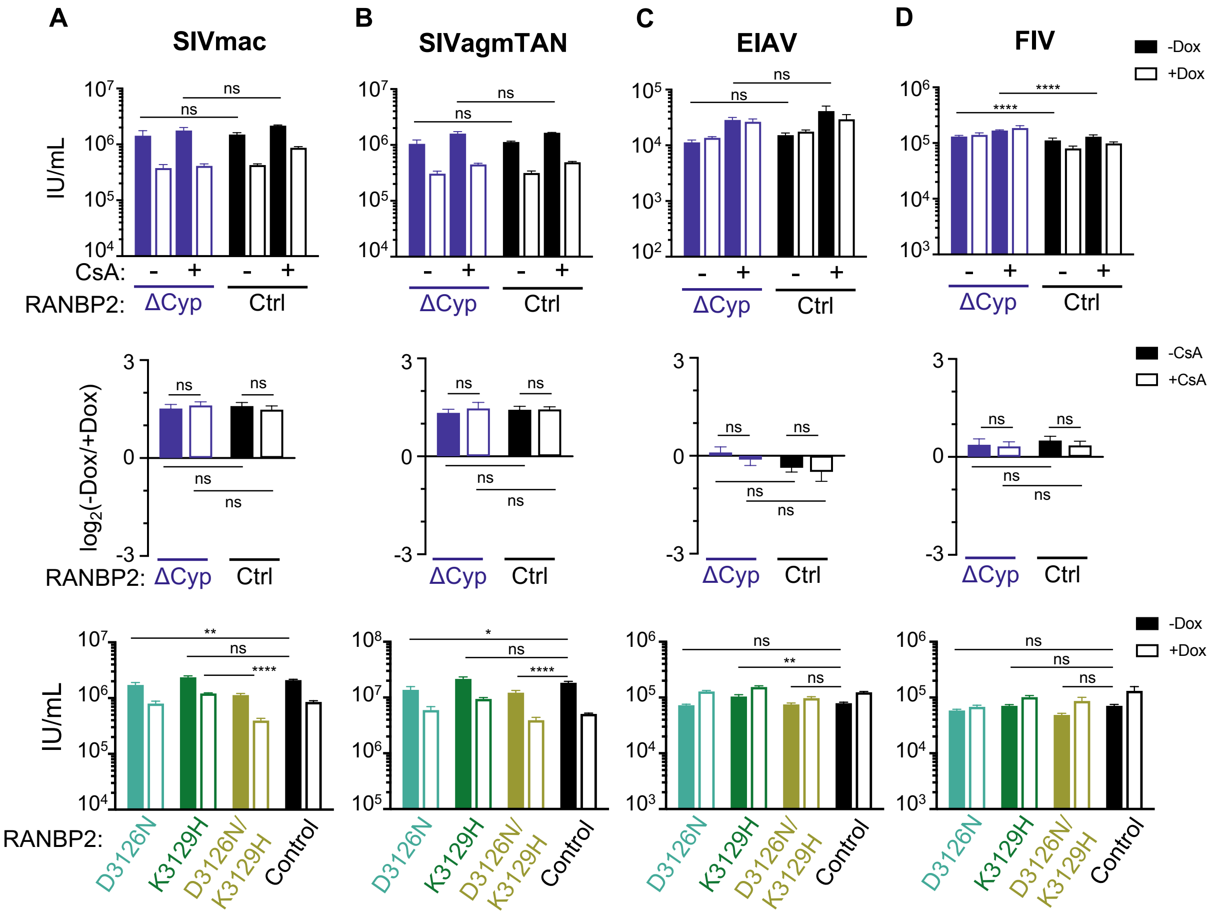


**Fig. S3. Effects of the RanBP2-Cyp domain on lentivirus infection and Mx2 sensitivity.**
A-D) (Top) Infection of control and RANBP2$_{\Delta Cyp}$ HT1080 cell clones stably transduced with doxycycline-inducible MX2 the presence (open bars) or absence (filled bars) of doxycycline and presence or absence of CsA with the indicated GFP reporter viruses. Titers are represented as mean + sem of infectious units (IU) per mL. RANBP2$_{\Delta Cyp}$: n≥12 technical replicates combined from ≥3 different clones; control: n≥8 technical replicates combined from ≥2 different clones; representative of ≥4 independent experiments. Statistical significance was determined by two-way ANOVA (Śídák's multiple comparisons test).
(Middle) Data from (top) shown as a ratio [fold change of -Dox(-MX2)/+Dox(+MX2)] in the presence (open bars) or absence (filled bars) of CsA. Average fold change calculated from three-four technical replicates per experiment; shown is mean + sem of $log_2$(fold change) from four-eight independent experiments. Statistical significance was determined by paired *t* test.
(Bottom) Infection of control and RANBP2 point mutant HT1080 cell clones stably transduced with doxycycline-inducible MX2 the presence (open bars) or absence (filled bars) of doxycycline with the indicated GFP reporter viruses. Titers are represented as mean + sem of infectious units (IU) per mL. n≥12 technical replicates combined from ≥4 independent experiments. Statistical significance was determined by two-way ANOVA (Śídák's multiple comparisons test)
ns, not significant (*P*≥0.05); * *P* < 0.05; ** *P* < 0.01; *** *P* < 0.001; **** *P* < 0.0001.

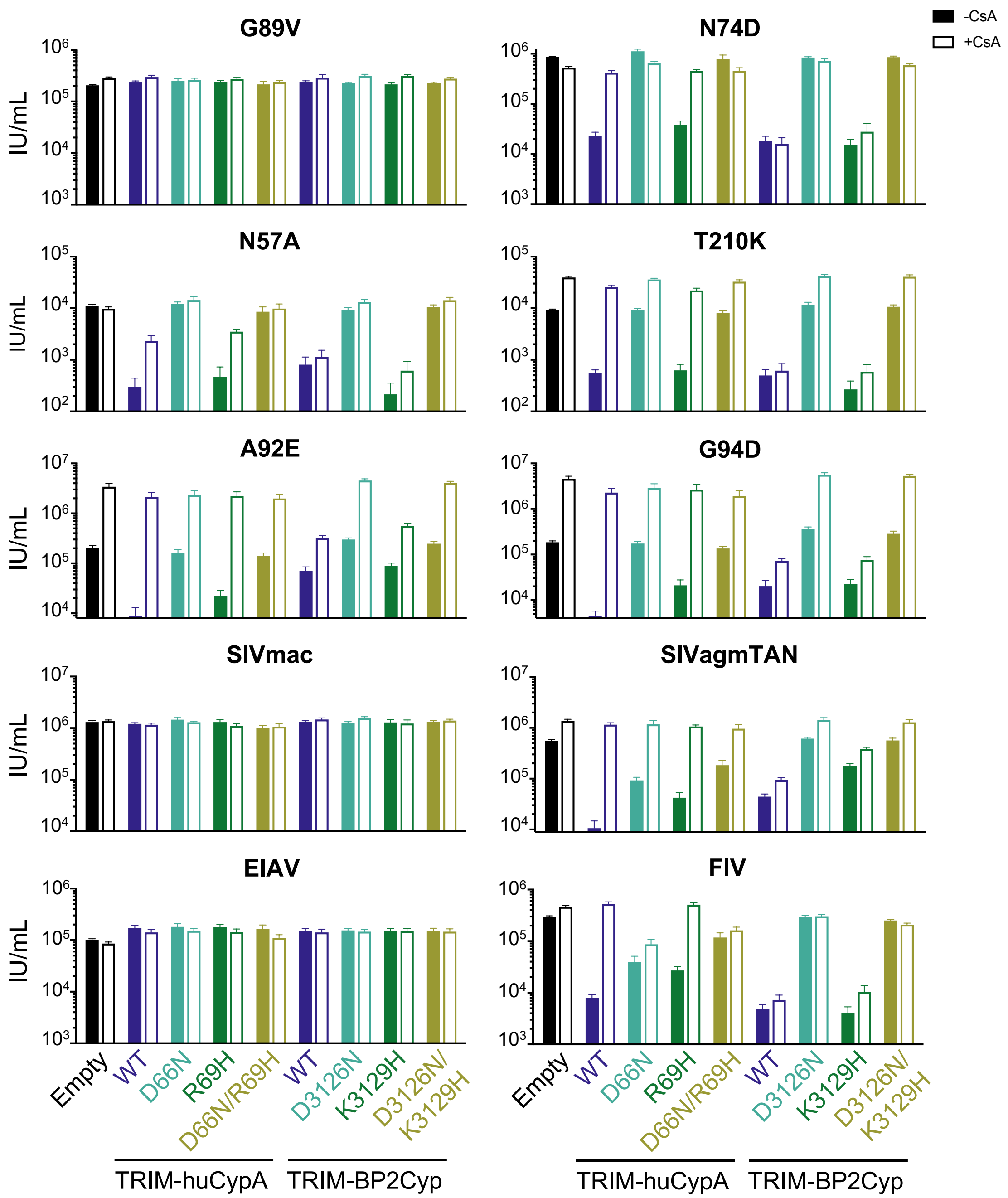


**Fig. S4. Recognition of lentiviruses and HIV-1 CA mutants by TRIM-Cyp fusions.**
Infectivity of GFP reporter viruses on HeLa cells stably expressing control empty vector, or chimeras of the TRIM5 N-terminal domain of owl monkey TRIMCyp with human cyclophilin A (huCypA), human $CypA_{D66N}$, $CypA_{R69H}$, $CypA_{D66N/R69H}$ mutants, human RANBP2-Cyp (BP2-Cyp), or $BP2\text{-}Cyp_{D3126N}$, $BP2\text{-}Cyp_{K3129H}$, $BP2\text{-}Cyp_{D3126N/K3129H}$ mutants. Titers are represented as mean + sem of infectious units (IU) per mL, n≥12 technical replicates combined from ≥3 independent experiments. Statistical analysis in Dataset S1.

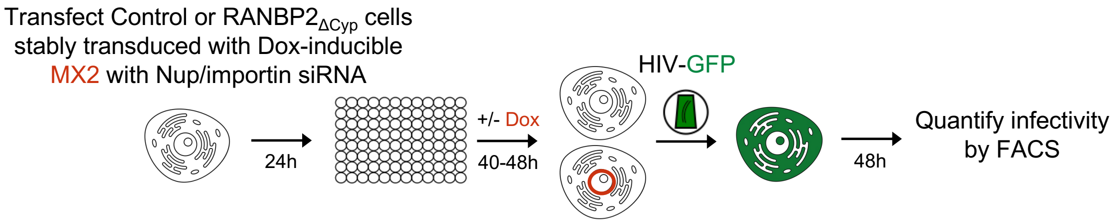


**Fig. S5. Experimental design for Nup/NTR knockdown**
Experimental strategy to investigate the roles of Nups and NTRs in HIV infection and the antiviral activity of MX2. For a detailed description, refer to the Methods.

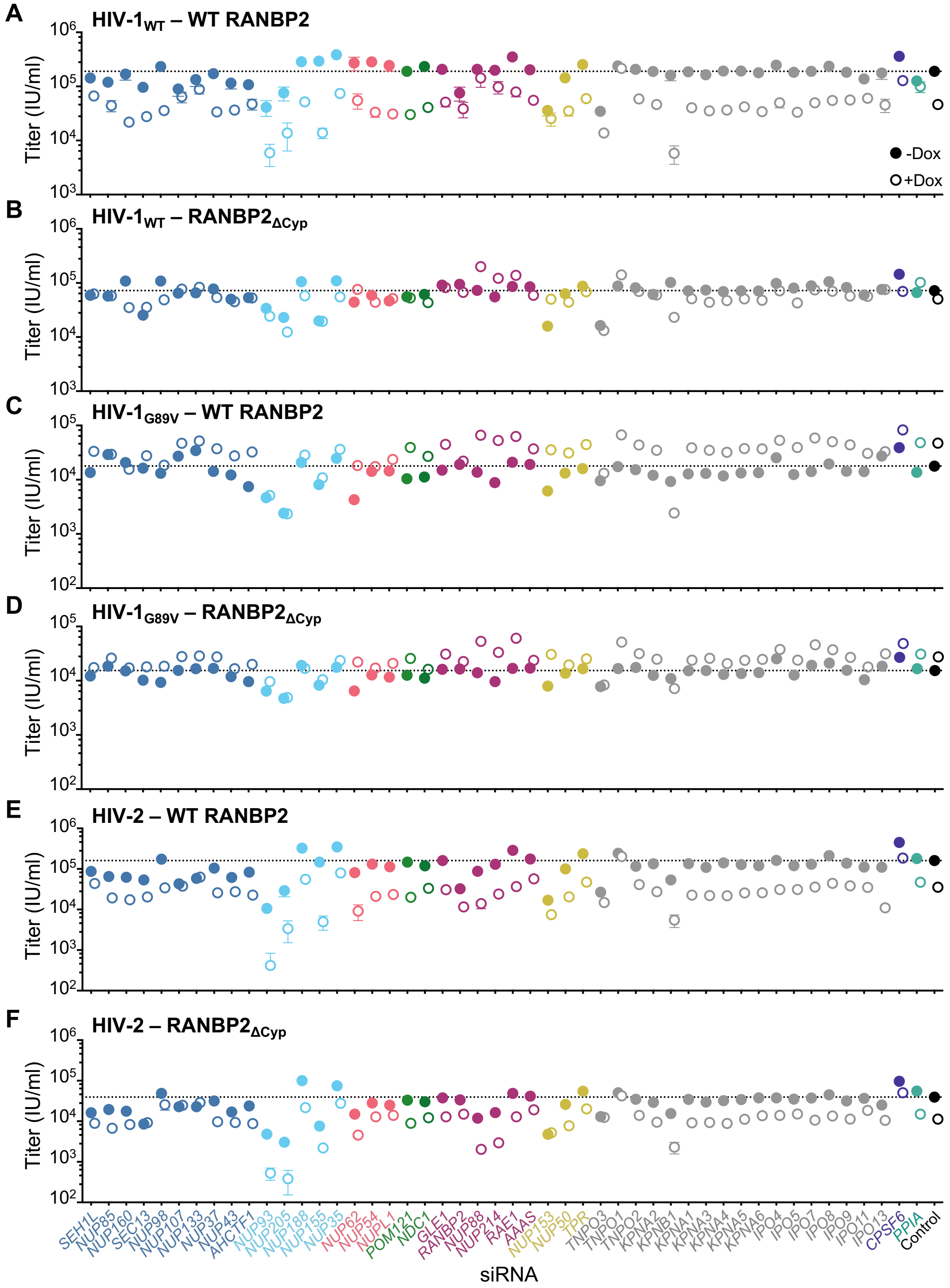


**Fig. S6. Effect of RANBP2-Cyp on the nucleoporin requirements for HIV infection and MX2 sensitivity.**

Infectivity of HIV GFP reporter virus in control (A,C,E) and RANBP2$_{\Delta Cyp}$ (B,D,F) HT1080 cells stably transduced with doxycycline-inducible MX2 in the presence (open circles) or absence (filled circles) of doxycycline 64-72h after transfection with siRNA (color coded by subcomplex as in Fig. 5A). Titers are mean ± sem, n=8 technical replicates from one control and one RANBP2$_{\Delta Cyp}$ clone from two independent experiments. Statistical analysis in Dataset S2.

### Table S1. Oligonucleotides for Plasmid Construction

| | |
|---|---|
| **TRIM5-fusions** | |
| BP2-Cyp NotI F | ataagaatgcggccgccAATCCTGTGGTGTTTTTTGATG |
| BP2-Cyp SalI R | ataagaatgtcgactcaTATCTGTCCACATTCTGTG |
| BP2-Cyp-DtoN-F | GTTTGCCAAGGAGGAAATATCACCAAACATGATGGAACAGG |
| BP2-Cyp-DtoN-R | CCTGTTCCATCATGTTTGGTGATATTTCCTCCTTGGCAAAC |
| BP2-Cyp-KtoH-F | GTTTGCCAAGGAGGAGATATCACCCATCATGATGGAACAGG |
| BP2-Cyp-KtoH-R | CCTGTTCCATCATGATGGGTGATATCTCCTCCTTGGCAAAC |
| BP2-Cyp-NH-F | GTTTGCCAAGGAGGAAATATCACCCATCATGATGGAACAGG |
| BP2-Cyp-NH-R | CCTGTTCCATCATGATGGGTGATATTTCCTCCTTGGCAAAC |
| **AAV** | |
| AAV NheI F | TTCCTGCGGCCGCAGCATAGCTAGC |
| AAV XhoI R | CGCTCGGTCCGCACAATTCCTCGAG |
| BP2-Cyp AAV KpnI F | AATACTAAGGTACCTGCTTCCCC |
| BP2-Cyp AAV NdeI R | TGAACAACATATGATTTTATATCTGTCCACATTC |
| **RANBP2-mutant cell screening and verification** | |
| BP2delCyp Check HR F | TGGAAGCTCTATTTATTAGATGTCTAGC |
| BP2delCyp Check HR R | CTTTGTAGTTCCATATTGAGTAGGCC |
| BP2 Exon 26 F1 | GAACCCAGTCAGCCGGTAAA |
| BP2Cyp R | AACACCACAGGATTGGTCTCC |
| BP2 Exon 28 F | ATGGAGAAGCAAAAGTAGAAC |
| Common Cyp R2 | TTTTCATCTTCAAATTTGTC |

### Table S2. ON-TARGET SMARTpool siRNA utilized in this investigation

| Gene Symbol | Gene ID |
|---|---|
| AAAS | 8086 |
| AHCTF1 | 25909 |
| CPSF6 | 11052 |
| GLE1 | 2733 |
| IPO11 | 51194 |
| IPO13 | 9670 |
| IPO4 | 79711 |
| IPO5 | 3843 |
| IPO7 | 10527 |
| IPO8 | 10526 |
| IPO9 | 55705 |
| KPNA1 | 3836 |
| KPNA2 | 3838 |
| KPNA3 | 3839 |
| KPNA4 | 3840 |
| KPNA5 | 3841 |
| KPNA6 | 23633 |
| KPNB1 | 3837 |
| NDC1 | 55706 |
| NUP107 | 57122 |
| NUP133 | 55746 |
| NUP153 | 9972 |
| NUP155 | 9631 |
| NUP160 | 23279 |
| NUP188 | 23511 |
| NUP205 | 23165 |
| NUP210 | 23225 |
| NUP214 | 8021 |
| NUP35 | 129401 |
| NUP37 | 79023 |
| NUP43 | 348995 |
| NUP50 | 10762 |
| NUP54 | 53371 |
| NUP62 | 23636 |
| NUP85 | 79902 |
| NUP88 | 4927 |
| NUP93 | 9688 |
| NUP98 | 4928 |
| NUPL1 | 9818 |
| POM121 | 9883 |
| PPIA | 5478 |
| RAE1 | 8480 |
| RANBP2 | 5903 |
| SEC13 | 6396 |
| SEH1L | 81929 |
| TNPO1 | 3842 |
| TNPO2 | 30000 |
| TNPO3 | 23534 |
| TPR | 7175 |

### Table S3. Antibodies utilized in this investigation

| Reactivity | Species | Company | Catalog Number |
|---|---|---|---|
| GAPDH | mouse | Santa Cruz | sc-47724 |
| HIV-1 p55+p24+p17 | rabbit | Abcam | ab63917 |
| MX2 | rabbit | Novus Biologicals | NBP1-8108 |
| RANBP2 | rabbit | Abcam | Ab64276 |
| NUP153 | mouse | Proteintech | Ab96462 |
| HA tag | mouse | BioLegend | 901514 |
| LAMIN B1 | Rabbit | Abcam | Ab133741 |
| Tubulin | mouse | Sigma-Aldrich | T6074 |

**Dataset S1 (separate file). Integration Site Analysis**
Raw data and oligonucleotide sequences for Fig. 4 are included in this file.

**Dataset S2 (separate file). Statistical Analysis**
Statistical analyses not included in figures (Figs. 3B, S4, and S6) is detailed in this file.
